# Universal abundance fluctuations across microbial communities, tropical forests, and urban populations

**DOI:** 10.1101/2022.09.14.508016

**Authors:** Ashish B. George, James O’Dwyer

## Abstract

The growth of complex populations, such as microbial communities, forests, and cities, occurs over vastly different spatial and temporal scales. Although research in different fields has developed detailed, system-specific models to understand each individual system, a unified analysis of different complex populations is lacking; such an analysis could deepen our understanding of each system and facilitate cross-pollination of tools and insights across fields. Here, for the first time we use a shared framework to analyze time-series data of the human gut microbiome, tropical forest, and urban employment. We demonstrate that a single, three-parameter model of stochastic population dynamics can reproduce the empirical distributions of population abundances and fluctuations in all three data sets. The three parameters characterizing a species measure its mean abundance, deterministic stability, and stochasticity. Our analysis reveals that, despite the vast differences in scale, all three systems occupy a similar region of parameter space when time is measured in generations. In other words, although the fluctuations observed in these systems may appear different, this difference is primarily due to the different physical timescales associated with each system. Further, we show that the distribution of temporal abundance fluctuations is described by just two parameters and derive a two-parameter functional form for abundance fluctuations to improve risk estimation and forecasting.

## INTRODUCTION

The dynamics of complex populations is studied in fields ranging from microbiology to economics. These studies have culminated in theoretical and computational models, with various levels of system-specific detail, that have made progress towards explaining system behavior and developing conceptual understanding. This includes the consumer-resource and Lotka-Volterra models of microbial communities; economic, econometric, and physics-based models of urban dynamics; and ecological models of forests based in niche and neutral theory^1–6^. Quantitative analysis, inspired by these models, has also unearthed statistical patterns in the data, hinting at simple emergent behavior of each population^7–11^.

What is lacking, however, is an investigation of emergent behavior using a common framework across different populations and an analysis of the temporal fluctuations in these populations. Classic models in statistical physics, such as diffusion and the Ornstein-Uhlenbeck process, have successfully described the behavior of diverse physical systems^12–14^. Their success demonstrates that some emergent properties are determined by only a few key underlying details of the dynamics. Efforts at applying this philosophy in other fields have sometimes succeeded^4,15–18^, but are often hindered by the lack of data and the incorporation of many system-specific details that make models difficult to analyze. In this paper, we undertake a statistical physics-inspired investigation of three different complex populations: microbial communities, tropical forests, and human cities, through a common framework. Our analysis will aim to uncover the key underlying similarities and differences between the populations; this will not only deepen our understanding of each system, but also facilitate the interconnection of tools and techniques between research fields.

All three complex populations we consider fluctuate in time. Large fluctuations in these populations are associated with catastrophic events such as disease, economic or ecological collapse^19–22^; hence understanding these fluctuations is crucial for risk assessment, quantitative biological methods, and forecasting^19,23–27^. Yet, many models of these populations study the equilibrium and steady-state properties such as the average population abundance ^1,2,22,28–31^. This restriction is in part because 1) analyzing dynamical properties of complex models is harder than analyzing their steady-state behavior and 2) temporal data required to fit and validate complex models has been lacking. We will tackle these two challenges by 1) adopting a minimal model which makes analysis of fluctuations feasible and 2) using data from three separate systems to fit and validate the model. Generating a reliable null model for population fluctuations that enables improved risk estimation is therefore a major goal of the study.

Fitting models using time series analysis has a precedent in each of the fields we consider here. For example, multiple studies have analyzed the possibility that simple models (making a range of assumptions) can reproduce empirical patterns in time-series data of gut microbiome communities^8,32,33^, tropical forests^34–37^, and urban populations^4,15^ (see SI Sec.4). Despite this history of previous analyses, we do not know of any comparison made across these data types using the same model, hence putting all three data sets on the same footing and amenable to direct comparison. We’ll thus go beyond earlier individual studies by analyzing both time-series and temporal snapshots of data simultaneously, using the same model applied across all three different systems, and introducing novel categorizations of each set of population data into distinct variant types motivated by domain-specific knowledge.

While these three systems span spatial scales from a single body to an entire country, temporal scales from days to years, and are studied in separate research fields, we harness the similarities in structure of the data to identify the same emergent features in all three systems. We compare the observed features to predictions from a model of stochastic population dynamics with just three population-specific parameters. This simplicity allows us to make analytical predictions for emergent features that we can fit and validate using the limited data. Remarkably, the model is able to capture most of the observed variation, despite its simplicity.

## MODEL AND DATA

To develop a unified understanding of complex population dynamics across spatio-temporal scales, we analyze time-series population data from three disparate systems: microbial communities in the human gut, trees in a tropical forest, and employment in U.S. cities (see Fig.1A). Traditionally studied by different fields, we interrogate the data using a shared framework by taking advantage of the similarities in the structure of the data. Each data set contains the relative abundance of population categories within each community at a sequence of time points (see Fig. 1B).

**FIG. 1.**
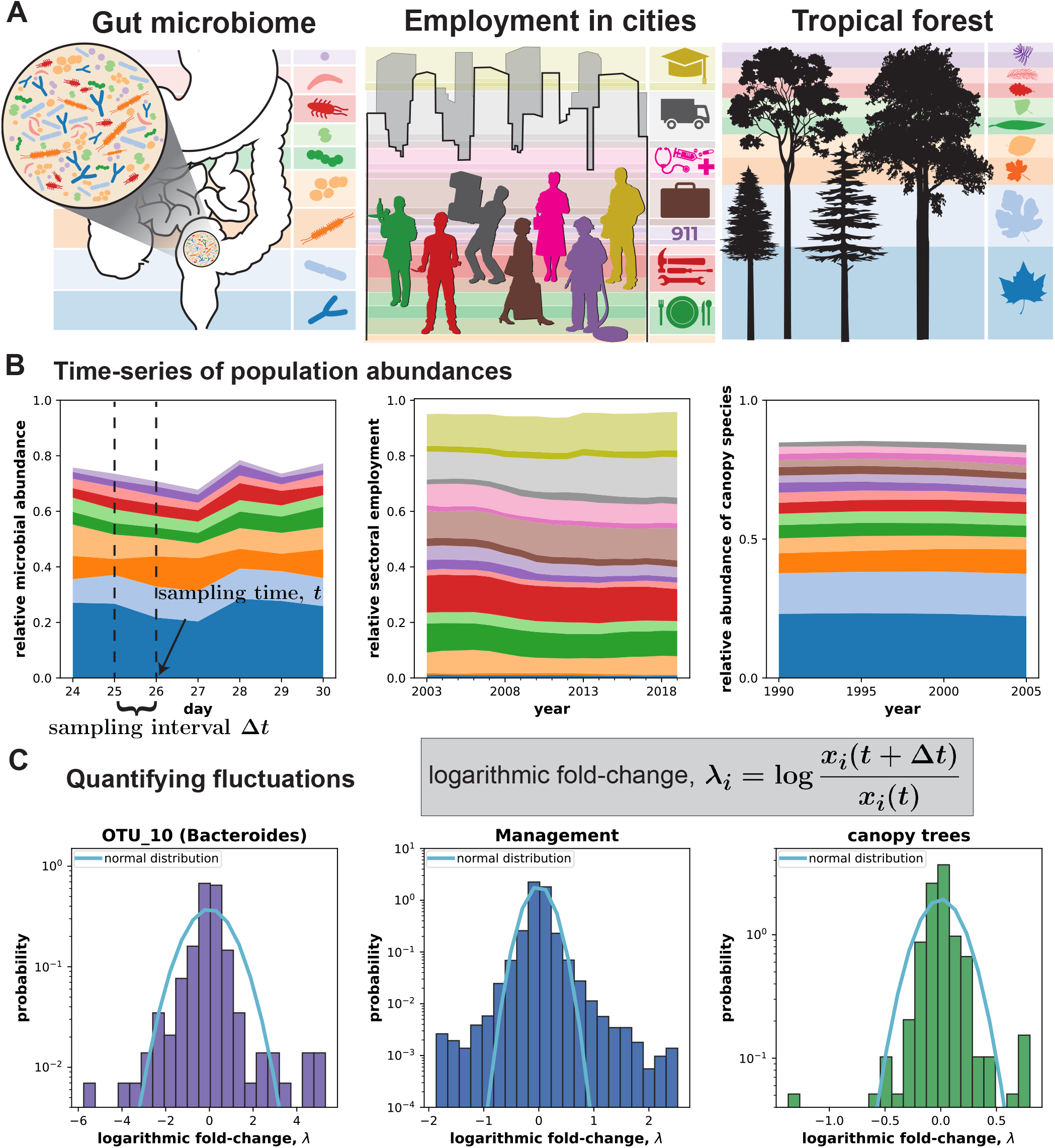
Time series data from three complex populations spanning spatio-temporal scales. **A)** We analyze time-series population data from three complex populations: the human gut microbiome, employment in U.S. cities, and trees in a tropical forest. Each complex population is composed of sub-populations, i.e., microbial species in the microbiome, employment within different economic sectors in cities, and tree species in four trait-based clusters in a tropical forest. **B)** Abundances of the sub-populations (species/sectors), sampled at periodic intervals, continue to fluctuate in time. Data shows microbiome abundance over a week, employment over 17 years for San Diego metropolitan area, and abundance of canopy tree species. **C)** The abundance fluctuations are measured using the logarithmic fold-change, 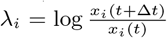, where *x*_*i*_ is the abundance of microbial species, an economic sector across cities, or tree species in a height cluster. The distribution of logarithmic fold change (LFD) for one microbial species, the management sector across all cities, and all canopy tree species is shown. The fit by the normal distribution to the LFD is shown by the cyan line. The fit is unable to capture the large fluctuations in the tails, central peak of the distribution, or both, illustrating that fluctuations in these complex populations are not normally distributed. Only the 10 (15) most abundant microbial (tree) species are shown in panel B for clarity.

More specifically, the gut microbiome dataset records the relative abundances of microbial species in the human gut sampled at daily intervals for almost a year^38^. A microbial species or Operational Taxonomic Unit (OTU) was defined based on genomic similarity^38^. The data from the Barro Colorado Island (BCI) forest records the number of tree species within a 50 hectare plot on the island, sampled at 5-year intervals for two decades^39^. We group trees based on clustering along trait axes into four clusters. There is a long history of grouping species by their maximum height^40^, based on the idea that species with access to similar levels, variability, or horizontal uniformity of light are more likely to compete strongly with each other^41^. Precisely which species to assign to which height cluster has since been put on a more quantitative footing^42^, with four distinct height clusters (shrubs, understory treelets, midstory trees, and canopy trees) identified on Barro Colorado Island. The employment dataset (https://www.bls.gov/cew/) records the number of people employed in different economic sectors, classified according to North American Industry Classification System (NAICS), in 383 US cities (Metropolitan Statistical Areas). The data is sampled monthly for 17 years.

We analyze the relative abundances in all three systems. Relative employment in a sector (employment in the sector in a given city, divided by total employment in the city) removes the effect of large variation in total population sizes of cities^5^. Similarly, we analyze the relative abundance of a tree species within a height cluster. Using relative abundances in both these systems allows us to treat data from all three systems on the same footing, normalizes out any temporal changes in the total population sizes stemming from overall population growth or decline (SI Fig.S10), and eliminates the effects of the large differences in city sizes. Our Methods section gives a mathematical definition of relative abundance in each system and SI Sec.8 and Figs.S10-S12 provide our analysis of absolute abundance data. Although working in terms of relative abundances introduces a constraint (that all relative abundances sum to equal one), for diverse communities this constraint has a relatively insignificant effect. We might also expect departures from our model for very high relative abundances, given that this constraint will tend to change the fluctuation properties for such variants as they approach the size of the entire system. But in practice, this is a small effect; it is rare in these data for any relative abundance to even approach 0.5. For the forest data, we show that our results are robust to the choice of relative vs absolute abundances by analyzing absolute abundances in SI Sec.8 and Figs.S11,S12.

In contrast to the static predictions from most deterministic models, the data in Fig.1B shows that population abundances continue to fluctuate in time. Furthermore, the strength of these fluctuations differs between the three systems with the largest fluctuations observed in the microbiome. To quantify the strength of fluctuations, we measured the logarithmic fold-change in abundance over a time interval ∆*t*, defined as

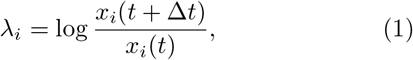

where *x*_*i*_(*t*) denotes the (relative) abundance of a species/sector *i* at time *t*. The empirical distribution of logarithmic fold-change or the Logarithmic Fold-change Distribution (LFD) is shown for a single species/sector from each system in Fig.1C. The comparison with the fit of the logarithmic fold-change by the normal distribution (equivalent to fitting fold-change by the lognormal distribution) illustrates that fluctuations cannot be understood as an outcome of environmental noise without any additional structure or mechanism (see Methods). Further, the fit by the normal distribution indicates that fluctuations much larger than expected from a normal distribution may occur in some of these complex populations. Understanding the distribution of abundance fluctuations is required to quantify the likelihood these large fluctuations which have a major impact on risk-estimates and time-series forecasting.

We now develop a simple model that is capable of predicting statistical features of all three data-sets, including the LFD. To achieve our goal of describing all three data sets, we keep the model both as generally-applicable and simply-formulated as possible. The model assumes that the abundance of a species/sector *i, x*_*i*_ fluctuates around an equilibrium value, 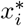, determined by the metabolic, ecological, or economic niche the species/sector occupies. We do not explore the system-specific mechanisms (resource competition, metabolic/economic interactions, etc.) that determine the particular equilibrium value^1,2,30,43–45^. Fluctuations then occur due to the stochastic processes governing population growth and decline. Deviations from the equilibrium value result in a linear restoring force, described by 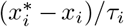, where *τ*_*i*_ is the timescale of return to equilibrium, where we neglect additional contributions from species interactions. Based on these assumptions, we call the model the Stochastic Linear-Response Model (SLRM) of population dynamics.

Assuming that population growth and decline occur in proportion to the abundance, we can write down the stochastic differential equation of the SLRM governing population abundances:

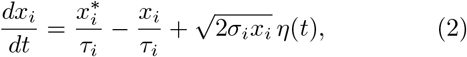

where *σ*_*i*_ captures the strength of population fluctuations, and *η*(*t*) is delta-correlated Gaussian noise or white noise. The model resembles the classic Ornstein-Uhlenbeck process, used to describe many stochastic quantities in physics and finance^12,14,46^, with one crucial difference: the scaling of the stochastic fluctuations with the square root of *x*_*i*_. This scaling reflects our assumption on the stochastic processes underlying population growth and decline; specifically, the square root scaling arises when growth and decline is a consequence of many independent, random events and the number of such events is proportional to *x*_*i*_, and is commonly used in many stochastic population models^47^. We note that this model has been proposed and analytical solutions were obtained previously in the context of singular diffusion processes by Feller^48^, bond interest rates by Cox and coauthors^49^, and in forest ecology (as a birth-death-immigration model) by Azaele et al^34^. Here, we go beyond these studies by applying a single model across three different systems. The SLRM has not to our knowledge been applied individually to employment data or microbiome data, and our application to forest ecology uses populations divided into novel, but ecologically-meaningful categories.

In ecological terms, the SLRM resembles a scenario where species occupy well-separated niches, with equilibrium abundance 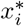 for species *i*. This idea of niche-separation determining model parameters is reflected in our assumptions. Specifically, in the microbiome, each species is described by its own SLRM parameters; in urban employment, each sector is described by its own SLRM parameters (independent of city); in forests, each trait-based cluster is described by its own SLRM parameters (independent of species index within the cluster). This assumption is reflected in our choice of showing the empirical LFD for a single sector across all cities, a single microbial species in the gut, and all tree species within a trait cluster in Fig.1C. While inter-species interactions may occur in each system, incorporating interactions drastically increases the model complexity hindering parameter-inference. Further, species interactions may not be necessary to describe population fluctuations in these systems, as we will show.

The minimal nature of the model allows us to make analytical predictions for two key quantities that characterize its long-term behavior. First, at long times (*t > τ*_*i*_ *>* 0) the distribution of population abundances will converge to a steady-state distribution. The steady-state distribution arises from the balance of stochastic fluctuations that kick the population from its deterministic equilibrium value and the restoring force towards the equilibrium. The analytical form of the distribution is (see Refs.^34,48,49^ and SI)

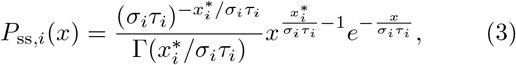

which is a Gamma distribution defined by two parameters: *σ*_*i*_*τ*_*i*_ and the Equilibrium to Noise Ratio (ENR), 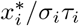, and Γ() is the gamma function. *σ*_*i*_*τ*_*i*_ quantifies the effective strength of the noise: i.e. the fluctuations over the equilibration time period. The ENR, 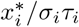, quantifies the relative strength of deterministic restoring force to stochastic fluctuations.

Second, we can analytically derive how abundances at steady-state fluctuate in time. Specifically, we quantify the temporal fluctuations by measuring the logarithmic fold-change of population abundances between two time points ∆*t* time units apart, denoted as 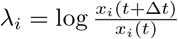 . The distribution of logarithmic fold-changes *λ*_*i*_ or Logarithmic Fold-change Distribution (LFD), in steady-state was derived in earlier studies (see Refs.^34,49^ and SI)

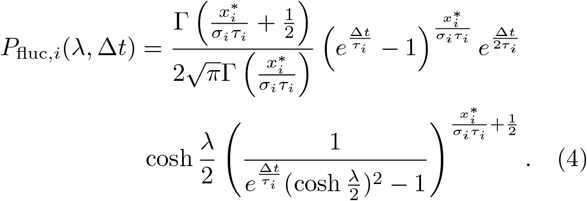

Despite its imposing appearance, the equation depends only on two parameters: the ENR, 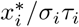, and the equilibration timescale, *τ*_*i*_, which scales the time interval ∆*t*. This distribution quantifies the likelihood of large abundance fluctuations in the model. In particular, the tails of the distribution at large *λ* decay exponentially at a rate proportional to the ENR, i.e, 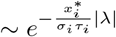, which ensures the moments of the distribution are well-defined. Note that this exponential decay for the logarithmic fold-change at large *λ* corresponds to a power-law decay for the fold-change, 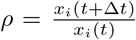, at large *ρ* with exponent 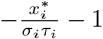.

## RESULTS

### Stochastic Linear Response Model (SLRM) reproduces empirical abundance fluctuations across populations

In line with our goal of characterizing abundance fluctuations in complex populations, we plot the empirical Logarithmic Fold-change Distribution (LFD) in the three systems in Fig. 2. Fig. 2A,B,C plots the LFD of urban employment, aggregated across all US cities, for three different sectors. Fig. 2D,E,F plots the LFD of 3 different species in the human gut microbiome. Fig. 2G,H plots the LFD of all the tree species belonging to two height cluster in the forest. The fold-change in abundance was calculated for time intervals of 1 year, 1 day, and 5 years for the city, microbiome, and forest data, respectively.

**FIG. 2.**
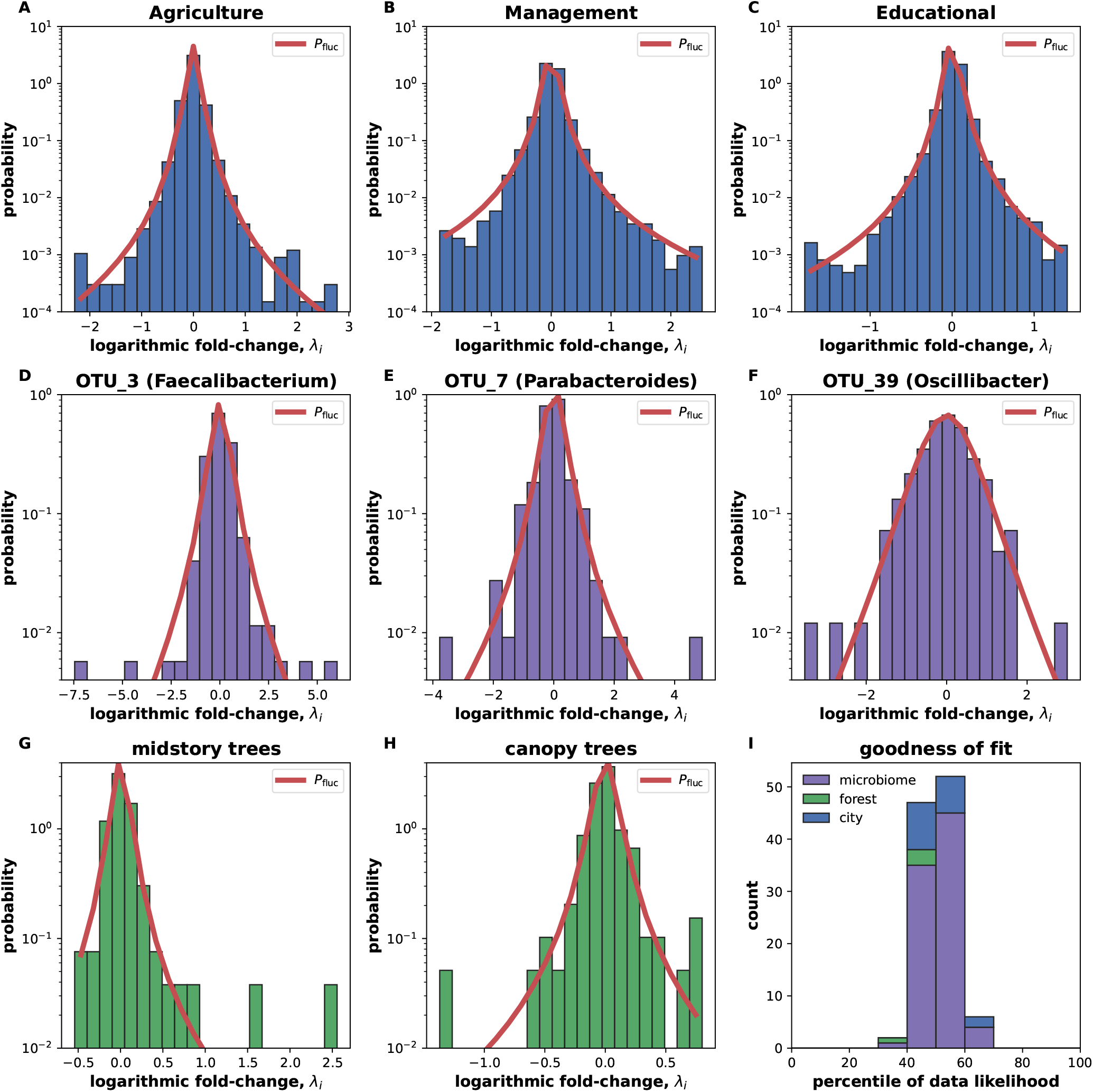
Universal distribution of fluctuations across complex populations. The histogram of the empirical Logarithmic Fold-change Distribution (LFD), i.e., the distribution of 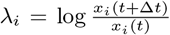, in each system is plotted and fit with the model prediction *P*_fluc_ from Eq. (4) (red line). The empirical distribution and fits are shown for: **(A-C)** employment in 3 different sectors aggregated across U.S. cities, **(D-F)** abundances of 3 microbial species in the human gut, and **(G**,**H)** abundances of tree species within two height clusters in the BCI forest. **I)** The percentile score, which quantifies the goodness of fit, shows that the likelihood of observing the data is similar to the likelihood of a random sample from the fitted distribution (of the same size as the data). Specifically, the percentile score quantifies the percentage of random samples with a higher likelihood than the observed data. A percentile score *>* 95% indicates a poor fit; all fits had a percentile score *<* 95%, as shown in the stacked histogram. The fold-change in abundance was calculated for time intervals of 1 year, 1 day, and 5 years for the city, microbiome, and forest data.

All LFDs were roughly centered around *λ*_*i*_ = 0 (average *λ*_*i*_ was 0.001, 0.004, and 0.05 for city, microbiome, and forest), but exhibited a large variation relative to the mean (coefficient of variation ≫ 1; see SI Fig.S1). This relative dominance of fluctuations over systematic trends for relative abundances not only makes the study of fluctuations easier, but also underscores the need to understand fluctuations better. The Stochastic Linear Response Model (SLRM), which neglects any systematic trends in the mean, is therefore a good candidate model of population fluctuations.

The red lines in Fig. 2 show the fit of the observed LFDs with the SLRM prediction, *P*_fluc_ (Eq. (4)). A separate fit is performed for each employment sector, microbial species, and forest niche, allowing the inference of corresponding population parameters. Despite having only two free parameters, the SLRM is able to fit the observed fluctuations across these diverse systems. We compared the SLRM fit to fits by two other candidate distributions using the Akaike Information Criterion^50^, as an additional test of the fit. The two candidate distributions (normal distribution and Laplace distribution) were chosen based on their success at fitting empirical fluctuations and alternative models (see Methods and Ref.^7,47,51^). The SLRM prediction outperformed the other candidate distributions in the majority of cases (see SI Fig.S2 and datasets 1,2,3). We note that our forest analysis demonstrates the importance of the distinct niches, given that inferred timescales range from 400 to 1700 years, and are significantly different between shrubs and other categories (see SI Sec. S7 and Fig.S13).

We then further quantified the fits, independent of comparisons with other models, by testing whether the observed data was likely to have been generated by the model. Specifically, for each fit and corresponding parameter estimates, we compared the likelihood of the observed data to the likelihood of 100 random samples of the same size as the data generated from the fitted distribution. This kind of ‘exact’ statistical significance test follows a precedent from testing neutral ecological models and population genetics models^52,53^. If the likelihood of the observed data was smaller than the 95% of the samples, we concluded that the data was unlikely to have been generated by the fitted distribution and rejected the fit. The percentile score in Fig.1I quantifies the percentage of random samples with a larger likelihood than the observed data. All fits had a percentile score *<*95%, meaning that SLRM passes our goodness of fit test in all cases. Thus *P*_fluc_ serves as a reasonable null expectation for the distribution of population fluctuations.

Comparing the three systems in Fig. 2, we notice that the scale of fold-change on the x-axis appears larger for the microbiome (Fig. 2D-F) than the two macroscopic systems (Fig. 2A-C,G-H). These differences in the data are reflected in the parameters values of the fit, and we explore this parameter variation in the next section. Importantly, despite these differences in scale and shape of the data, *P*_fluc_ provides a good fit to data from all three systems.

### Universal fluctuations appear different in micro- and macroscopic systems due to different generation times

Fitting *P*_fluc_ to the empirical LFD provides us with maximum likelihood estimates for the equilibration timescale *τ*_*i*_ and the ENR 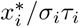. We compare the dimensionless versions of these inferred parameters across the three systems. Specifically, we compare the ratio of the equilibration timescale to time interval of observation *τ/t*_*obs*_ and the ENR across the three systems.

Fig 3A plots the ENR and *τ/t*_*obs*_ across the three systems. While the ENR spans a similar range across all three data sets, the micro- and macroscopic systems differ in the inferred values of *τ/t*_*obs*_. For microbial populations, *τ/t*_*obs*_ *<* 1, which means that the duration over which we observe the system, about a year, is longer than its equilibration timescale which is around 10 days. In contrast, cities and forests have *τ/t*_*obs*_ *>* 1, implying that the duration of observation, two decades, is much shorter than the equilibration timescale. This difference in *τ/t*_*obs*_ is responsible for the difference in shapes of the fitted *P*_fluc_ in Fig. 2. Thus, the duration of observation is an important difference between microscopic (microbiome) and macroscopic systems (cities and forests).

**FIG. 3.**
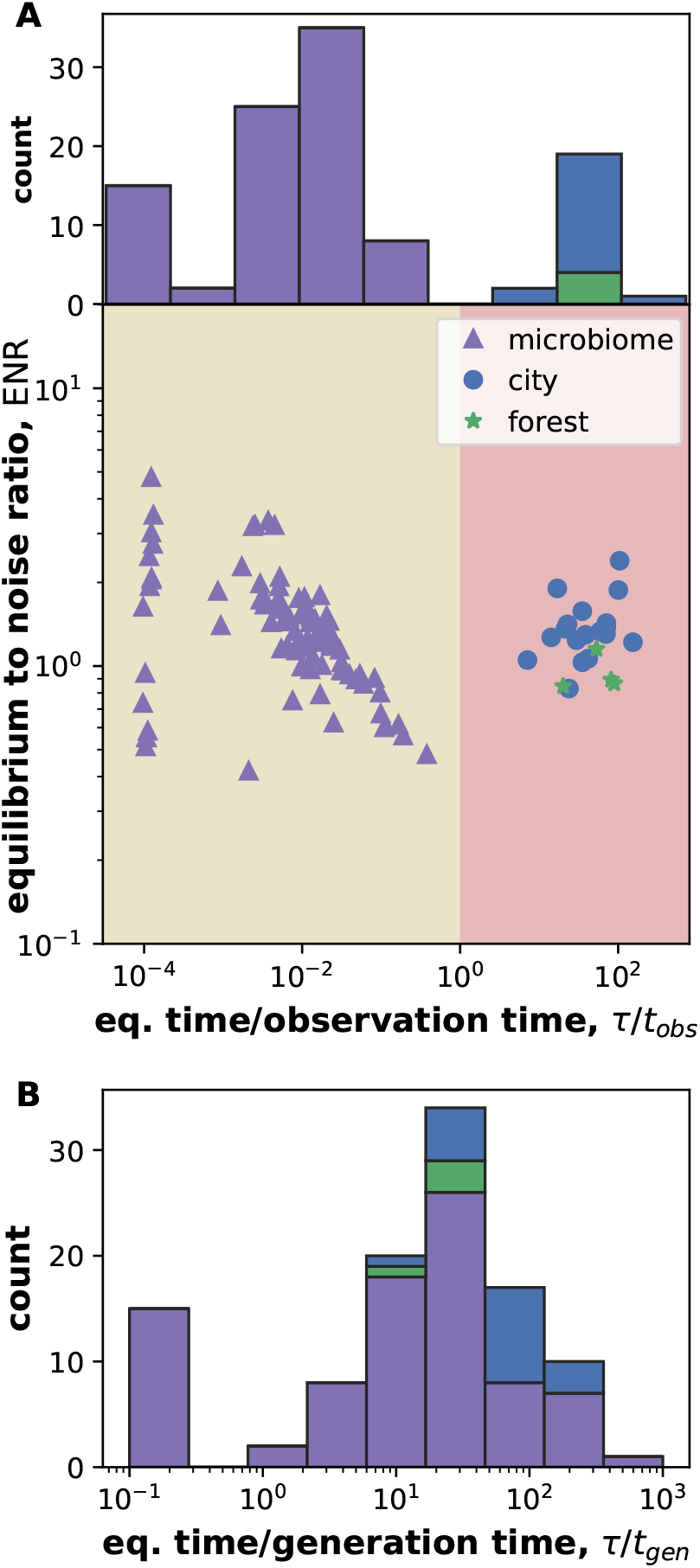
Timescale of observation differentiates micro-scopic and macroscopic systems; generation time unifies them. **A)** Plot of the dimensionless parameters ENR 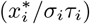 and *τ/t*_*obs*_ inferred from fitting the empirical LFD. *τ/t*_*obs*_ is the ratio of the equilibration timescale to time duration of observation. All three systems span similar values in ENR, but *τ/t*_*obs*_ differentiates the microscopic and macroscopic systems. Microbiome data has *τ/t*_*obs*_ *<* 1, i.e., we observe the microbiome for longer than its equilibration times. The macroscopic systems have *τ/t*_*obs*_ *>* 1, i.e., we observe cities and forests for shorter than their equilibration times. **B)** Measuring time in generations reveals the similarities between systems. The stacked histogram of inferred *τ/t*_*gen*_, the ratio of the equilibration timescale to the generation time, shows that all 3 systems inhabit a similar parameter range, with an equilibration timescale on the order of 10 generations. A small group of microbes appear as outliers with *τ/t*_*gen*_ *<* 1; these are the microbes for which the LFD was well-fit by a normal distribution (see SI Fig.S3). The estimated generation time, based on Refs.^54–57^, was 4.2 hours, 10 years, and 55 years for microbiome, cities, and forests (see Methods). See SI Table S1 for the full name of each sector.

An alternative way to compare these populations is to use the ratio of the equilibration timescale to the generation time, *τ/t*_*gen*_ (see Fig 3B). The generation times in the 3 systems were estimated as 4.2 hours, 10 years, and 55 years for microbes, employment, and forests respectively (see Methods and Refs.^54–57^). Remarkably, when viewed in terms of generation time, all 3 systems occupy a similar region of parameter space. Hence, fluctuations in the three systems are described by the same distribution over a similar parameter range when time is measured in generations, highlighting the similarities in the emergent behavior across the population types. We note that a small subset of microbes in Fig 3B appear as outliers, with *τ/t*_*gen*_ *<* 1. For these microbes, the empirical LFD is better-fit by a normal distribution than the model prediction *P*_fluc_ (see SI Fig.S3). For the remaining data, the equilibration time scale is on the order of 10 generations for all three systems. Therefore, when viewed in terms of generation time rather than physical time, emergent fluctuations in the three systems are highly similar.

In addition to examining how the inferred timescale, *τ*, varied across the three systems, we also examined how *τ* varied within each system. Analyzing employment data, we found that the most abundant sectors in cities such as healthcare and retail trade had the longest timescales while the least abundant sectors in cities such as agriculture and mining had the shortest timescales. Quantitatively, we found that the median relative abundance of a sector across cities was correlated with the inferred timescale (Pearson’s r =0.86, Spearman r =0.66, *p <* 0.05). We found a similar relationship for microbes, with the most abundant species (belonging to Bacteroides) having the longest timescales (Pearson’s r =0.93, Spearman r =0.68, *p <* 0.05) (SI Fig.S5). For forest clusters, we found that shrubs had a significantly shorter timescale than the other three, taller height clusters (see SI Sec.7, Figs.S13). We will return to interpret these correlations shortly.

### Stochastic Linear Response Model reproduces empirical distribution of species abundances

From Fig. 3, we see that the observation duration for the microbiome is longer than its equilibration time scale (*τ/t*_*obs*_ *<* 1). Hence the temporal trajectory of abundances of each microbial species should trace out the corresponding steady-state distribution, which is a gamma distribution described by two parameters ((3)). We plot the distribution of relative of microbial species and fit it with the two-parameter gamma distribution (black line), via maximum likelihood in Fig.4A,B,C.

For city and forest data, the observation timescale is longer than the equilibration timescale (*τ/t*_*obs*_ *>* 1), and so the temporal trajectory of abundances will not converge to the steady-state distribution. Instead, we plot the cross-sectional distribution of abundances, i.e., the relative abundance of a sector in a city aggregated across all cities and the abundances of tree species within a height cluster, at a random time point (Fig. 4D-H). The fit with the gamma distribution is the black line.

**FIG. 4.**
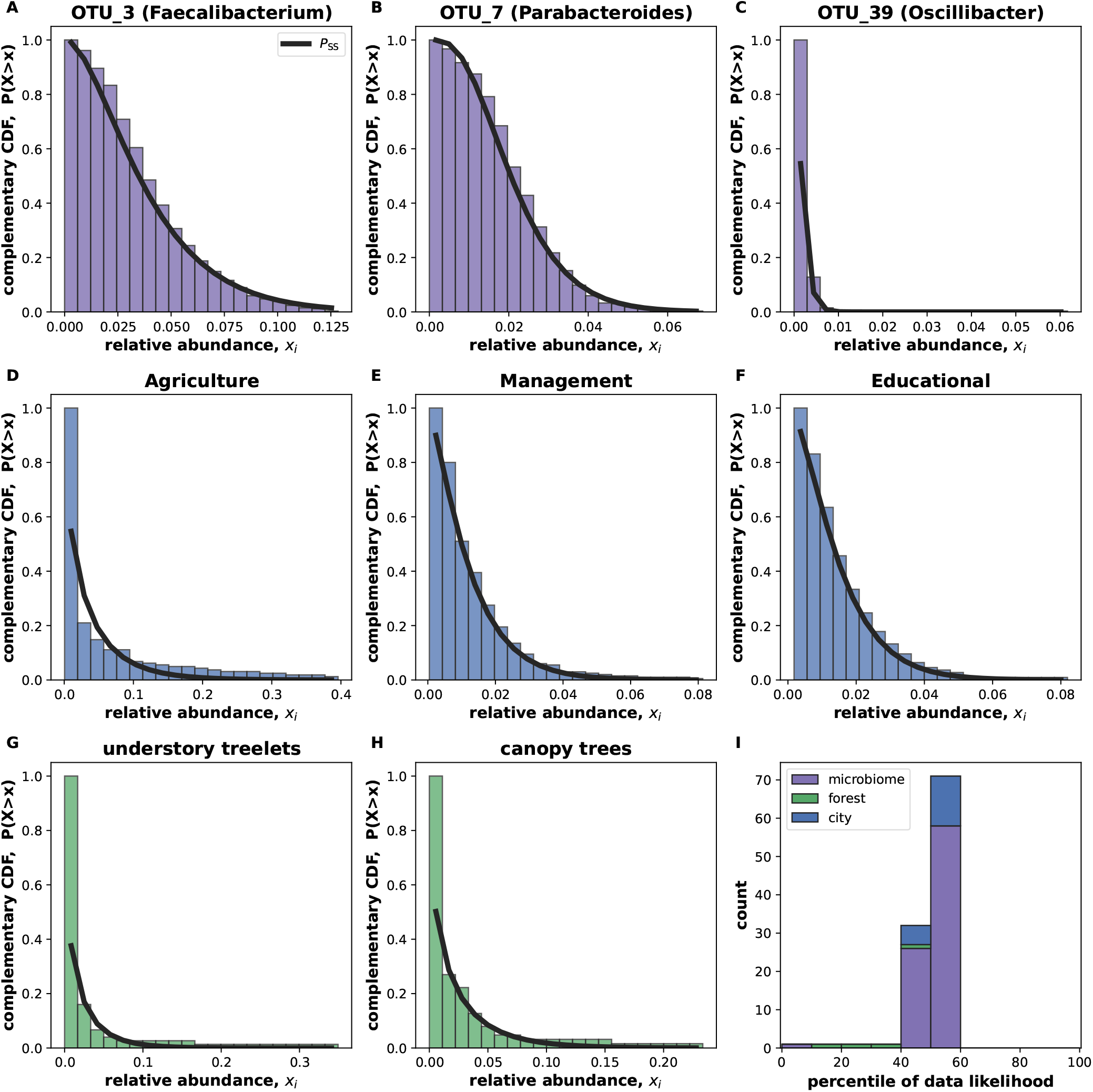
The distribution of abundances is well-fit by the SLRM steady-state distribution. **A-C)** The distribution of microbial abundances over time for three different species is fit by the two-parameter gamma distribution predicted by the SLRM (black line). We fit the temporal distribution of microbial abundances because we observe the microbiome for far longer than its equilibration time scale. **D-H)** For cities and forests, we observe the system for intervals shorter than the equilibration time. Hence, we fit the cross-sectional distribution of abundances, i.e, employment in a sector across all cities and abundance of all tree species in a height cluster. Data and fit for the three species, three sectors and two height clusters are shown as the complementary cumulative distribution function 1 − *CDF* (*x*) i.e., the probability that a values i greater than the x-axis *P* (*X > x*). **I)** The percentile score, which quantifies the goodness of fit, shows that the likelihood of observing the data from the fitted distribution is similar to the likelihood of a random sample of the same size as the data. All fits had a percentile score *<* 95%, as seen from the stacked histogram. Two additional goodness of fit tests were also performed and the majority of the species pass both tests (see SI Fig.S6)

To test whether the observed data could have been generated by the model, we repeat the procedure used in Fig. 2. Specifically, we compared the likelihood of the observed data to the likelihood of randomly generated samples from the fitted distribution. All data sets had a likelihood comparable to a random sample, as evidenced by the percentile scores (from 1000 random samples of the same size) shown in Fig.4I. Two additional goodness of fit tests were also performed and the majority of species passed both tests (see SI Sec.S5 and Figs. S6, S7). Together, these observations indicate that the abundances in these systems are well described by the gamma distribution. This conclusion is further supported by previous research that used a gamma distribution to successfully fit cross-sectional microbial abundances across microbiomes^8^.

The steady-state distribution (SSD) and logarithmic fold-change distribution (LFD) predicted by the model ((3), (4)) share a parameter, the ENR 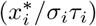. We compare the maximum likelihood estimates of the ENR obtained by the separate fits of the empirical abundance distribution and LFD in Fig. 5. The discrepancy in inferred ENR values could arise from limitations of the data or the model.

**FIG. 5.**
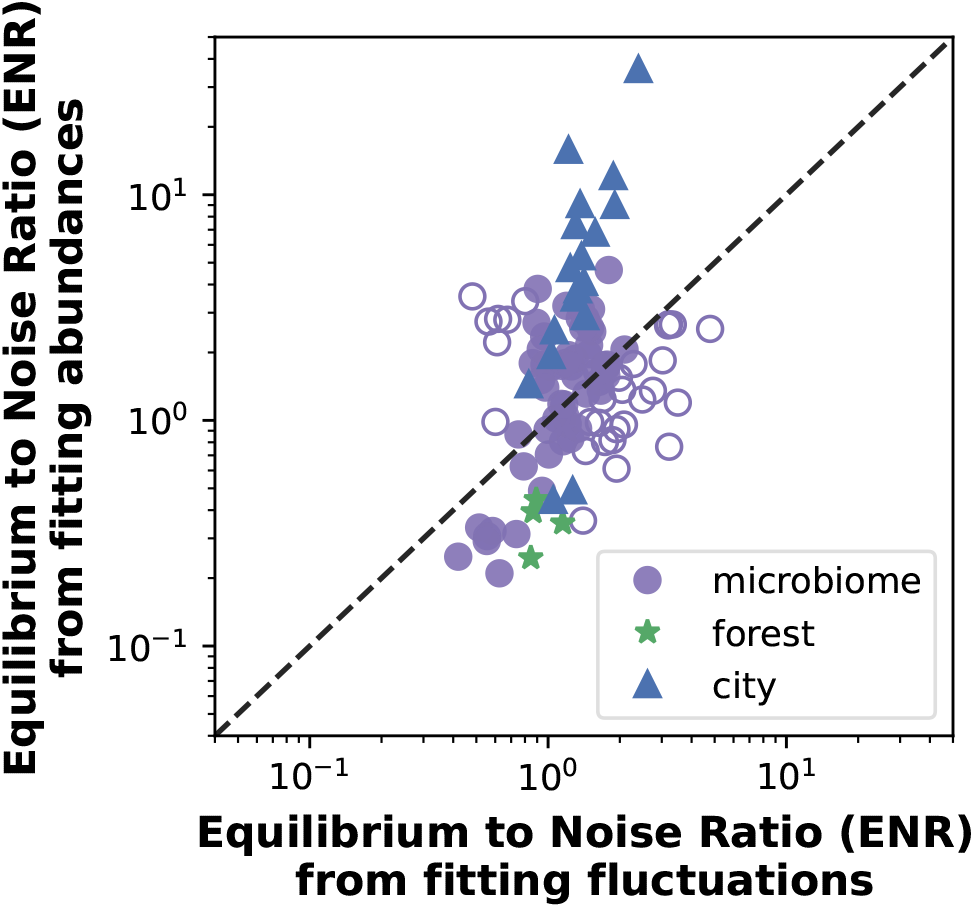
The consistency of inferred parameters from independent fits of the distributions of fluctuation and abundance. Independent estimates of the Equilibrium to Noise Ratio 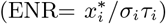 can be inferred from fitting the empirical Logarithmic Fold-change Distribution (LFD) or ftting the abundance distribution. If a fit of the LFD using the ENR value inferred from fitting the abundance passes the goodness of fit test, or vice-versa, we deem the inferred values of ENR as consistent. Filled points denote consistent estimates and unfilled points denote inconsistent data points. All city sectors and forest species were consistent, and 57 of 85 microbial species were consistent.

To check whether the model is able to simultaneously fit both the LFD and abundance distribution, we performed a modified version of our goodness of fit test we used previously. First, using the ENR inferred from fitting the abundance, we compare the likelihood of fitting the LFD by *P*_fluc_ to the likelihood of 100 random samples from the fitted distribution of the same size. Then we fit the other way around, i.e., using the ENR inferred from fitting the LFD, we compare the likelihood fitting the abundance by *P*_ss_ to the likelihood of 1000 random samples from the fitted distribution of the same size. If the likelihood of the observed data was within 95% of the likelihood of the samples for either of these comparisons, then we conclude that the inferred parameters were consistent, i.e., the ENR obtained from fitting one distribution is able to provide a reasonable fit of the other distribution. The data points deemed consistent are depicted by filled markers in Fig. 5. If the data likelihood was smaller than 95% in both cases, we term the inferred parameters as inconsistent. The data points deemed inconsistent are shown as unfilled markers in Fig. 5.

The ENR estimates for the majority of the data were consistent (Fig. 5). A minority of the microbes (28/85) were rejected as being inconsistent. This could partly be due to temporal correlations in the data used to fit the abundance distribution, which we neglected. These correlations exist over timescales of *τ*_*i*_ and only vanish when ∆t ≫*τ*_*i*_. For cities and forests, we used cross-sectional data, which does not suffer from this drawback. All employment data, including the apparently large outliers in Fig. 5, were consistent. The consistency of these large outliers was because although the abundance distribution of some sectors were well fit by large ENR values, substantially smaller ENR values also provided a reasonable fit and could not be rejected using the few hundred observations. Overall, the majority of the observed species/sectors in the three systems followed the expected relationship between the predicted distributions for abundances and fluctuations.

### Variation of model parameters explains Taylor’s law

In addition to providing a simple 2-parameter null model for fluctuations and abundances in complex populations, the SLRM can also help understand other empirical patterns in the data. Prior research in microbiome data has revealed an approximate power-law scaling (with exponent 2) of the variance vs. mean of the abundance, called Taylor’s law^7,8^ (Fig. 6), which may arise due to various mechanisms^58–60^. Examination of inferred model parameters provides an alternative explanation for this empirical observation.

**FIG. 6.**
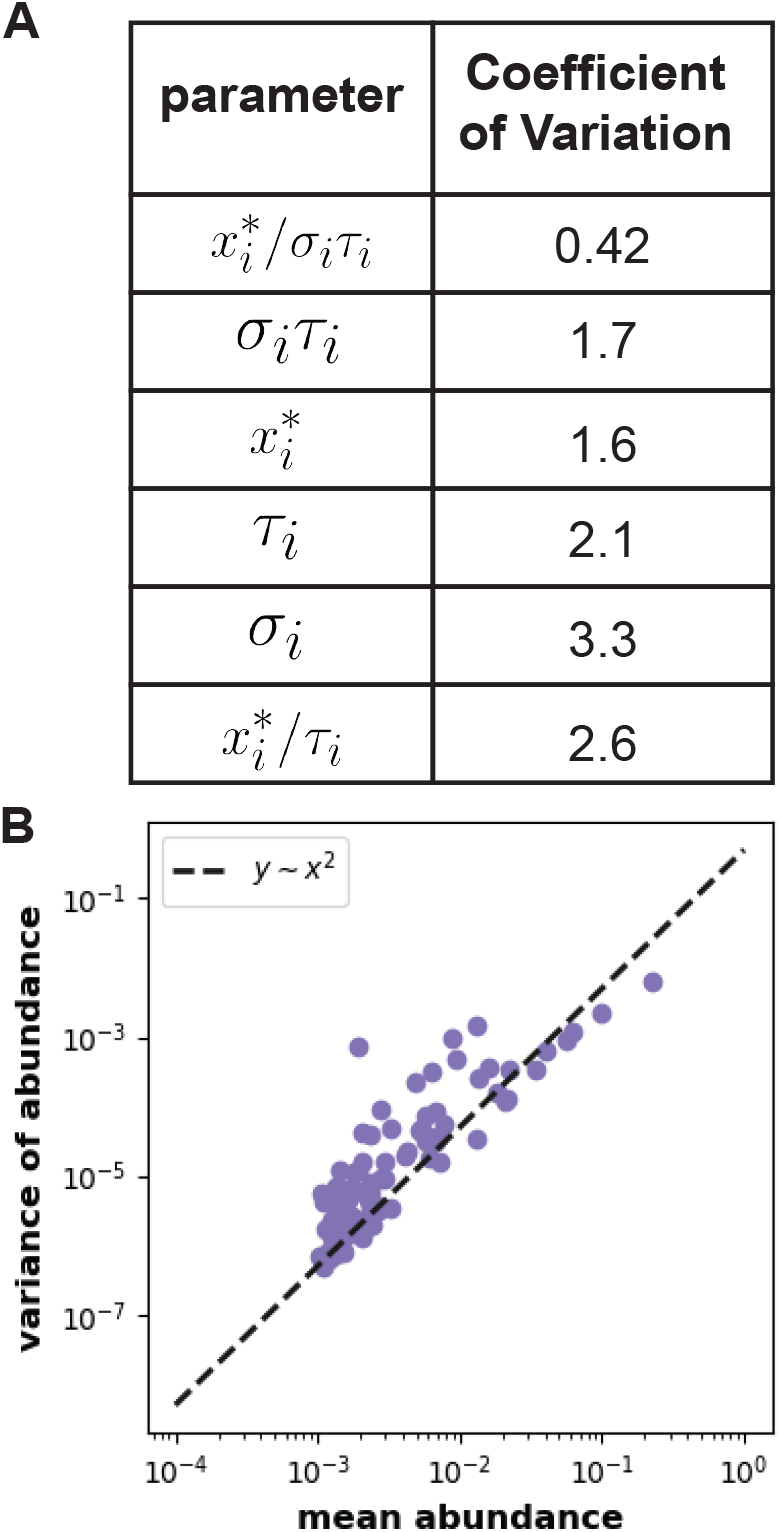
Power-law scaling of temporal mean and variance of abundance (Taylor’s law) in microbiome due to approximate conservation of ENR. **A)**The inferred ENR 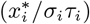 in the microbiome data is relatively constant compared to other parameters, as evidenced by the tabulated coefficient of variation. **B)** The properties of the gamma distribution (which fits the abundance distribution) define the mean abundance of species *i*, as 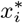, and the ratio of variance to mean squared as the ENR 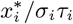. Thus, due to the approximate conservation of the ENR, we get the power-law scaling (with exponent 2) of variance with mean referred to as Taylor’s law. The geometric mean of the ENR inferred from fitting the LFD and abundance distribution was used when estimating the variation of different parameter combinations.

To investigate why Taylor’s law arises, we compare the variation of the inferred model parameters across microbial species, using the coefficient of variation (Fig. 6A). The ENR 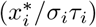 has a substantially lower coefficient of variation than the other parameters, and so can be considered to be approximately constant. For the Gamma distribution, the mean abundance of a species *i* is 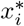 and the ratio of variance to mean squared is the ENR, 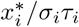. In microbiome data, the ENR is constant, and so the ratio of variance to mean squared remains fixed while the mean abundance varies, which leads to the observation of an approximate power-law scaling of the variance with mean. Hence, the approximate constancy of the ENR provides an alternative explanation for the observed power-law known as Taylor’s law in microbiome data. The approximate constancy of ENR implies that the distribution of fluctuations (LFD) of different microbial species will be similar when time intervals are measured in terms of *τ*_*i*_. Furthermore, the approximate constancy may explain one of our earlier findings— that relative abundance correlates with inferred timescale (SI Fig.S5). Mathematically, since the ENR does not vary significantly, relative abundance in our model must be proportional to *σ*_*i*_*τ*_*i*_. Future work may uncover the mechanisms behind why ENR is approximately constant across these systems.

### Comparing SLRM and a model with environmental noise

The Stochastic Linear Response Model (SLRM) incorporates ‘square-root’ fluctuations, referred to as demographic noise, and commonly used in many population dynamics models^47^. Demographic noise captures the fluctuations arising from accumulation of small, independent random growth and death events, and mathematically it can be identified by the square-root scaling of the noise term with population size (Eq. 2). An alternative form of noise used in population dynamics models is environmental noise^47^. Environmental noise aims to capture fluctuations arising from random fluctuations of the overall growth and death rates of the population, and has been used in the analysis of both microbiome data^8^ and local forest communities^35–37^. The latter involve a range of choices of model specification, including the way competition is imposed in the local community, and how dispersal is modeled from regional pool to local patches. This complexity tends to yield models without analytical solutions for LFD and SSD. On the other hand, a relatively simple implementation of environmental stochasticity is the Stochastic Logistic Model (SLM), which has been applied to recapitulate the observed abundance distributions in another of our three data types: microbiome communities^8,32^. All of these models are characterized by the same linear scaling of the noise term in the population size (Eq. 5), and so comparing our model with an environmental noise model provides an initial test of whether environmental stochasticity will inevitably tend to provide a better description of fluctuations than demographic noise alone. Therefore, in this section we compare the SLRM, which incorporates demographic noise, with the SLM.

The SLM is also a three-parameter model defined by the following equation for the relative species abundance *x*_*i*_:

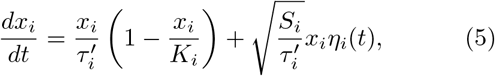

with parameters *K*_*i*_, which describes the carrying capacity of the population, *S*_*i*_, which captures the strength of fluctuations, and 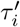, which sets the timescale of growth and equilibration. *η*_*i*_ is delta-correlated Gaussian noise or white noise.

The Steady-State abundance Distribution(SSD) of the SLM is also a Gamma distribution^8^, like the SLRM:

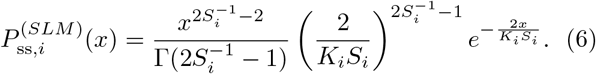

It is parameterized by combinations of the two parameters *K*_*i*_ and *S*_*i*_. However, unlike the SLRM, there is no analytical prediction for the Logarithmic Fold-change Distribution of the SLM.

Therefore, to facilitate a direct comparison between the SLRM and SLM, we adopted the following procedure: first, we fixed two of the three parameters in both models by fitting the gamma-distributed SSD predicted by the models to the observed abundance distribution. Then, we simulated the SLM for a range of values of remaining parameter, *τ* ^*′*^, to obtain a series of predicted LFDs from SLM simulations. We obtained the LFD for the same range of *τ* values of the SLRM through the analytical predictions. Finally, we compared the disagreement between the two sets of predicted LFDs and the empirical LFDs in the three systems by computing the Jensen-Shannon Distance between them (see Methods for further details).

In Fig. 7A,B, we illustrate this procedure applied to data on the employment in the management sector in US cities. The distributions predicted by the two models as the timescale parameters (*τ* ^*′*^, *τ*) are varied is shown by the colored lines alongside the observed data (black circles). The disagreement between the model prediction and the observed data is measured using the Jensen-Shannon Distance (JSD) and shown in the insets. The model with a lower JSD better explains the observed data. We repeated this analysis for all microbial species, employment sectors and forest niches, and found that the SLRM provided a better fit in the majority of the cases (Fig. 7C). Specifically, the SLRM had a lower JSD in 17 out of the 18 employment sectors, 72 out of 85 microbial species, and 2 out of the 4 forest clusters. Interestingly, it is the forest data, where a body of evidence exists to demonstrate the importance of environmental noise in explaining other aspects of population fluctuations^61,62^, where the SLRM and SLM were most similar in their performance. But outside of these cases, the SLRM provides the better description of the data.

**FIG. 7.**
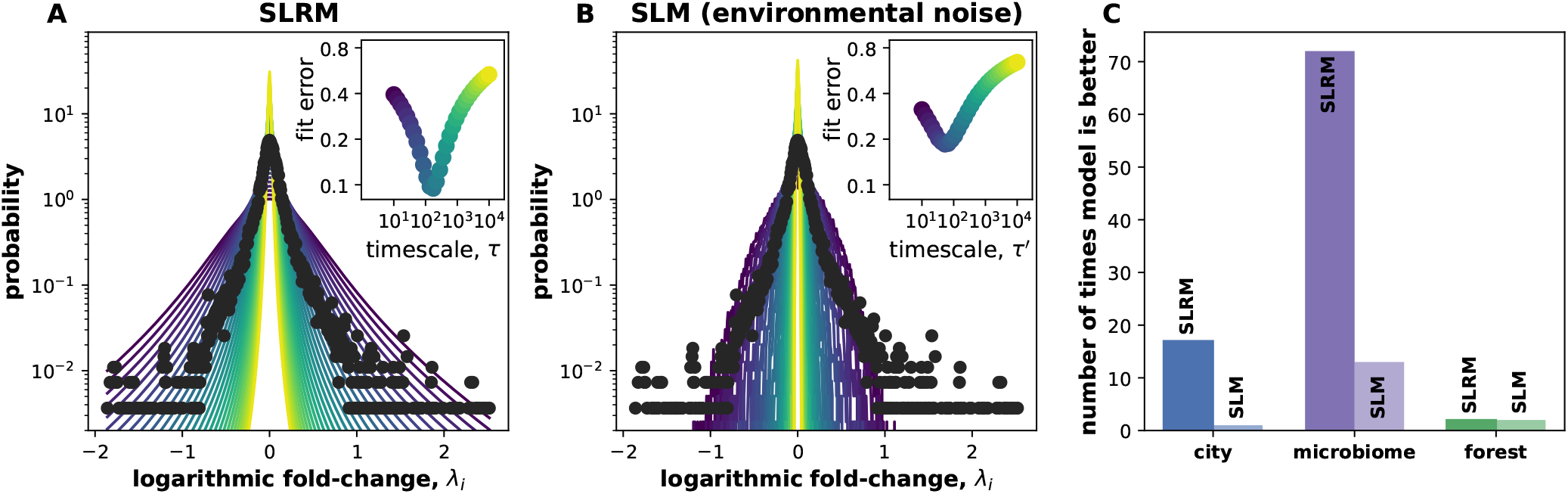
Comparing SLRM and a model with environmental noise (SLM). **A**,**B)** We compare the predicted Logarithmic Fold-change Distribution (LFD) of the SLRM, which incorporates demographic noise, and the SLM, which incorporates environmental noise, to the empirical LFD (black points). Colored lines in panels A and B show the LFD of the SLRM and SLM varies for the same range of parameter (*τ* ^*′*^, *τ*) values; empirical data corresponds to employment in the management sector. The insets show the error between the model predictions and observed data, measured using the Jensen-Shannon Distance. A lower value of the JSD indicates a better fit; the SLRM fit this data better than the SLM. **C)** We repeated this procedure for all sectors, species and clusters to identify the model that provided a better fit (lower JSD) in each case. The SLRM provided a better fit to majority of the data—in 17 out of the 18 employment sectors, 72 out of 85 microbial species, and 2 out of the 4 forest clusters.

## DISCUSSION

Understanding the behavior of diverse complex populations is challenging, but critical for progress in many fields. For the first time, we analyze all three of microbiome data, forest data, and employment data on an equal footing. This unified approach to studying the three populations complements detailed investigations of each specific population. In the microbiome context, where many studies focus on species interactions^63,64^, our work demonstrates how statistical features of the data can arise from stochastic fluctuation, with minimal interactions. For forest data, our work finds some independent support for prior analysis that groups trees into distinct height niches^42^. And for employment data, where many studies have focused on detailed econometric models and analyses^6,43,65^, our work provides a novel perspective from the point of view of a simple dynamical model. Moreover, by showing that the emergent distributions of population abundances and fluctuations in all three populations can be reproduced by a single model, our analysis highlights emergent properties that are independent of system-specific details. When using generations to measure time, all three systems occupy a similar range of parameters, suggesting that fluctuations in cities and forests over decades and centuries would closely resemble microbiome fluctuations. In broader terms, the fact that our analysis provides a good description across a variety of systems may point to the existence of universal properties in the fluctuations of complex systems.

In quantitative terms, our analysis provides a simple two-parameter functional form for the distribution of fold-changes in the populations and relates this to the abundance distribution. The predicted distribution of fluctuations *P*_fluc_ ((4)) is able to fit fluctuations in all three systems despite having only two parameters, and moreover we find that when measured in terms of generation time, parameter fits for all three systems collapse into a narrow window of fitted values. Further, *P*_fluc_ arises from a well-defined model and suggests a plausible mechanism; it is not simply chosen from the vast library of statistical distributions historically examined. The observation that we find all three data types are described by similar parameter values suggests important, deeper connections that may yet be uncovered in future work.

The predicted distribution of fluctuations *P*_fluc_ can serve as an important null model of the fluctuations in complex populations, where understanding the likelihood of large fluctuations is crucial. For employment fluctuations, large fluctuations impact urban planning and economic stability; for forests, large fluctuations impact ecological management strategies; for microbiome, large fluctuations can cause dysbiosis, which affects the health of the host^19,20,22^. A two-parameter null model for fluctuations in complex populations is useful in practical, data-limited settings; it can estimate the risk of large fluctuations more accurately (SI Fig.S4) and improve quantitative methods, such as those utilising Bayesian inference from time-series data to classify ecosystem states^66^ and priors for priors for decision-making and modeling^67–69^. In the SI, we show how *P*_fluc_ fits empirical data from the three systems when fluctuations are measured over different time intervals (see SI Sec.S6, Figs. S8,S9).

The Stochastic Linear Response Model (SLRM) can be understood as the linearization of a more complex non-linear model around its equilibrium when species interactions are neglected. This is shown in our SI, where we also demonstrate (in SI Sec.S2) an example where a model with inter-species interactions actually reduces exactly to the SLRM. In general though, this simplification drastically reduces the number of parameters, making parameter inference from available data feasible. Deviations from the model predictions, could indicate the presence of species interactions, which are often modeled by Lotka-Volterra, consumer-resource and other models of higher order species interactions^1,2,63^, autocorrelated noise, or other mechanisms. Such models could potentially be parameterized by using specialized methods with additional data^25,70^. In the SI, we discuss how the SLRM can be extended into a stochastic model incorporating species interactions in a linear regime. Analysis of this extended model could pave the way for novel inference methods that account for the stochastic fluctuations in observational data.

While we compared our model predictions with a range of classic distributions, we also note that environmental noise^47^ has been proposed as an explanation for fluctuations in abundance across different complex systems, including multiple forest data sets^35–37^ and microbiome data^8^. To capture this mechanism and compare our model to its predictions, we tested the performance of the SLRM to the Stochastic Logistic Model (SLM)^8,32^, a three-parameter model that combines nonlinear logistic growth with environmental stochasticity. The SLM lacks an analytical solution for the Logarithmic Fold-change Distribution, but through numerical simulations we compared the fits of the SLRM and SLM to the empirical data. We found that the SLRM outperformed the model with enviromental noise in the majority of our data (Fig. 7). While there are multiple other types of enviromental noise model, for example those that have provided a good description of local forest community fluctuations^36,37^, this comparison demonstrates that environmental stochasticity does not necessarily provide a better description of fluctuations in complex populations. More general models of environmental stochasticity tend to lack simultaneous analytical solutions for the Logarthmic Fold-change Distributions and Steady-State Abundance distribution, making numerical comparison more challenging. However, it is certainly possible that, just as with species interactions, more general kinds of noise should form part of the basis for extending our model, and future analysis will likely shed light on this question.

Framing the SLRM as a useful base model for further research, we note that it can be easily augmented with additional mechanisms, including environmental fluctuations and species interactions, which could be tested with additional data (see SI). Other modifications could help understand evolving populations. For the timescales examined, we assumed that the equilibrium value *x*^∗^ remains constant. Over longer timescales, however, the equilibrium value could change due to biological evolution, climate change, or socio-technological revolution. Investigating the model when *x*^∗^ changes in time could help understand emergent dynamics in complex populations over evolutionary timescales and presents an interesting direction for future research.

To butcher two well-worn phrases, all models are wrong, some are useful, and some are unreasonably effective. We believe the SLRM falls into the latter two categories, and that its surprising effectiveness across such diverse datasets points to something universal about the way complex populations fluctuate. The SLRM also provides valuable two parameter null models for the distributions of abundances and fluctuations in complex populations, which are of particularly utility in data-limited scenarios for forecasting and risk-analysis. The unified analysis of the three population types highlights both similarities and differences between the systems, and paves the way for a fruitful exchange of tools, techniques, and interpretations between these very different fields.

## METHODS

### Data processing

#### City data

Public domain city data were obtained from Quarterly Census of Employment and Wages from the U.S. Bureau of Labor Statistics (https://www.bls.gov/cew/). The data provides the employment classified into industrial sectors by the North American Industry Classification System (NAICS) at county level in the U.S. (see SI Table S1). We aggregated data at the county level to 383 Metropolitan Statistical Areas (MSA), which we call cities. MSAs are independent statistical units defined by the Census Bureau that encompass a central city and the geographical areas connected to the city. We obtained the list of counties in each MSA in 2017 from U.S. Census Bureau, County Business Patterns program, and used this to calculate employment at MSA level. This approach allowed us to maintain a consistent definition of MSAs across the entire time-series. The employment data is recorded at monthly intervals from 2003 to 2019. Since many industries, such as agriculture and accommodation, display seasonal trends in employment withing a year, we used a ∆*t* of 1 year for calculating the empirical LFD. The duration of observation, *t*_*obs*_, was 17 years for employment data. We plot and fit data with only non-zero abundance.

For privacy reasons, sectoral employment data at some points are suppressed, and so we remove these points from our analysis. Since this suppression increases at finer levels of NAICS classification, we analyzed sectors classified at the two digit level. This also made our analysis robust to changes in the NAICS classification scheme that impacted the temporal continuity of the data at finer resolution. We used 18 of the 21 NAICS categories at the two-digit level (see SI Table S1). We removed three categories: ‘81’ (other),’92’ (public administration) and ‘99’ (unclassified). Public administration and government employment was removed because much of this sector is not reported due to governmental regulations. Relative sectoral employment was calculated for each city by dividing sectoral employment in the city with the total employment reported in the city. Note that due to data suppression and removal of three naics categories, the sum of the relative employment in a city over all sectors we analyze need not equal one. We assume that each sectors is described by a set of SLRM parameters that is independent of the city; this provides us with enough data to fit the SLRM predictions.

#### Microbiome data

Microbiome data from Ref.^38^ was obtained and processed as in Ref.^7^. We consider only the gut microbiome data for individual M3 since it was substantially longer than other time-series. The data was collected in time intervals of one day with some gaps in sampling, and hence ∆*t* = 1 day for microbiome data. There were a total of 336 time-points recorded excluding sampling gaps. Since there were sampling gaps, an approximate duration of observation, *t*_*obs*_, of 300 days was used for microbiome data.

To process the data, first, read counts at each time point were normalized to obtain relative abundance. Only prevalent or relatively abundant species were used for our analysis. Specifically, we considered species only if they were present in more than half of the time points and their average abundance was greater than 10^−3^. 85 species that met this criterion. To calculate the empirical LFD, we used abundances that were collected ∆*t* = 1 days apart. We plot and fit data with non-zero relative abundance. Each species has its own set of SLRM parameters.

#### Forest data

The Barro-Colorado Island forest data was obtained from the Center for Tropical Forest Science website (https://forestgeo.si.edu) ^39^. Abundance data collected at 5 year intervals, in years 1990, 1995, 2000, and 2005 was used. Only trees that were alive and had a diameter at breast height *>* 10 cm were counted. The trees were grouped into four height clusters: shrubs, understory treelets, midstory trees, and canopy trees based on Ref.^42^. There were 87, 75, 60, and 63 species in the four height clusters. Relative abundance of a species in a particular height cluster was calculated as the absolute abundance of the trees of the species at that time point divided by the absolute abundance of all tree trees within that specific height cluster. We assume that all species within a height cluster and thus being highly similar in trait values, are described the same set of SLRM parameters; this provides us the data required to fit the data with the model predictions. Different height clusters are fit separately, like different employment sectors. Time interval for calculating LFD, ∆*t*, was 5 years. The duration of observation, *t*_*obs*_, was 20 years for forest data.

### Fitting and sampling procedures

The data was fit and parameters were estimated by Maximum Likelihood Estimation from the Scipy package in Python. In addition to the probability distribution *P*_fluc_, the corresponding cumulative distribution (see SI) was defined to make sampling from the distribution more efficient.

In the city data set, we had substantially more data points in the each LFD (∼30, 000 points) than for the empirical abundance distribution (∼300 points). This made performing the goodness of fit test computationally harder for the LFD than the abundance distribution, since the test required generationg samples from the distribution of the same size as the data. Hence for each fit of the abundance distribution, we obtained 1000 samples to compare with the data likelihood, whereas for each fit of the LFD, we obtained only 100 samples.

The coefficient of variation of tabulated in Fig. 6A, used inferred parameters values of ENR, *σ*_*i*_*τ*_*i*_, and *τ*_*i*_ to construct the various parameter combinations. *τ*_*i*_ was estimated from fitting the LFD, *σ*_*i*_*τ*_*i*_ was estimated from fitting the abundance distribution, and the ENR was the geometric mean of the estimates from the LFD and abundance distribution.

### Definition of relative abundance in each system

For microbiome data, the relative abundance of species *i* at time *t, x*_*i*_(*t*) was defined as

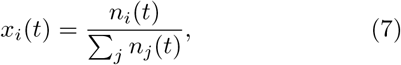

where *n*_*i*_(*t*) is the number of sequence read counts of the species obtained at time *t*. Since the number of read counts is different from the species number, we do not know the absolute abundance of microbes in the data. For the forest data, relative abundance of species *i* in cluster *c* at time *t, x*_*i*_(*t*), is given by

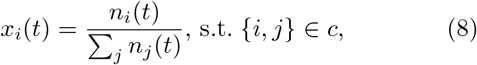

where *n*_*i*_(*t*), *n*_*j*_(*t*) are the counts of species *i, j* belonging to the same cluster *c* at time *t*. Hence, species abundance is normalized by the total population within the same height cluster to obtain its relative abundance. Eq. 8 can also be used to define relative abundance in city data. For city data, *x*_*i*_(*t*) is the relative abundance of sector *i* in city *c* at time *t* and *n*_*i*_(*t*), *n*_*j*_(*t*) is the employment in sectors *i, j* in the same city *c* at time *t*. Hence, sectoral employment is normalized by the total employment in the same city to obtain its relative abundance. This allows us to neglect the wide variation in total population sizes across cities^5^.

### Estimating generation times in each system

The generation times used for scaling the inferred time scale in Fig. 3 were calculated based on estimates in the literature. Since a direct estimate of generation time of microbes in the human gut is unavailable, we used the measured generation time in mice on a regular diet of divisions per day(4.2 hours)^54^. For trees, the generation time used was 55.5 years^55^. For urban employment, we measured generation time as the time required for people to change jobs between industrial sectors. Using the median job duration in US of 5 years^56^ and the fact that roughly half of the job changes are between sectors^57^, we obtained a generation time of 10 years. The estimated generation times neglect possible variation between species and sector due to data limitations.

### Simulating a model with environmental noise (SLM)

We implemented the Stochastic Logistic Model (SLM) using a temporal finite difference scheme in python. To ensure accurate simulation results, we chose time steps that were sufficiently small. Instances of the noise, *η* were generated by sampling a normal distribution with variance scaling appropriately with the time-step. To avoid species extinction, we imposed a small minimum population size. We simulated the population for longer than 10*τ* ^*′*^ to ensure that dynamics reached the steady-state. We then computed the LFD from the second half of the simulated data.

To compare the fits to the data of the SLRM and SLM, the same number (30) of logarithmically-spaced values of *τ* ^*′*^ (SLM) and *τ* (SLRM) were scanned for each system, and the JSD between the empirical and model LFDs were computed. The particular range of *τ* ^*′*^ and *τ* values scanned differed between the three systems; they were selected to ensure that a minima in JSD existed between the bounding values. The bounding values of this interval were 10, 10^4^ years for cities; 0.08, 100 days for the microbiome; and 2.5, *×* 5 10^4^ years for forests. We computed the JSD on binned data when comparing the LFD of the SLRM and SLM with the observed data. The binning was determined using the Freedman-Diaconis estimator on the observed data. Note that since we do not have an analytical prediction for the SLM, we cannot directly compute the likelihood of the observed data to measure a. The JSD, on the other hand, can be computed between the binned observed and predicted distributions.

### Environmental stochasticity can produce normally distributed fluctuations

The dynamics of a population driven purely by environmental noise is given by

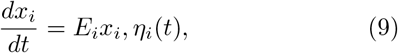

where *E*_*i*_ quantifies the strength of environmental noise and *η*_*i*_ is delta correlated Gaussian noise or white noise^47,61^. We can rewrite this equation for log *x*_*i*_ instead as

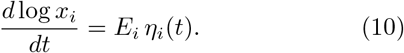

Clearly, the fluctuations of log *x*_*i*_ are now normally distributed. Thus environmental noise can produce a normally distributed LFD. Note, however, that this equation does not have a steady-state. Additional terms are required in the dynamical equation to stabilize the population and ensure a steady-state.

Although a subset of microbes have an LFD that is well-explained by a normal distribution, we are able to fit the majority of the data without using environmental noise. While adding environmental noise would affect model behavior, the success of the model without environmental noise suggests that the populations could be in a parameter-range where the effect of environmental noise is unimportant or that the quantities we examine are not sensitive to the addition of environmental noise on top of demographic noise.

## Data availability

All datasets analyzed in this manuscript are publicly available. Code used to process and analyze the data as described is available on Github (https://github.com/ashish-b-george/Universal-fluctuations)^71^. Employment data at the county level was obtained from the Quarterly Census of Employment and Wages from the U.S. Bureau of Labor Statistics (https://www.bls.gov/cew/downloadable-data-files.htm). Microbiome data from Ref.^38^ was obtained and processed as in Ref.^7^. Forest data was obtained from the Center for Tropical Forest Science website^39^.

## Acknowledgements

The authors would like to thank Zachary Miller, Alice Doucet Beaupré, and members of the O’Dwyer group for helpful feedback and comments. The authors acknowledge funding support from Simons Foundation Grant #376199 and McDonnell Foundation Grant #220020439 to J.O.D. The BCI forest dynamics research project was made possible by NSF grants to S.P. Hubbell: DEB #0640386, DEB #0425651, DEB #0346488, DEB #0129874, DEB #00753102, DEB #9909347, DEB #9615226, DEB #9405933, DEB #9221033, DEB #9100058, DEB #8906869, DEB #8605042, DEB #8206992, DEB #7922197, support from CTFS, the Smithsonian Tropical Research Institute, the John D. and Catherine T. MacArthur Foundation, the Mellon Foundation, the Small World Institute Fund, and numerous private individuals, and through the hard work of over 100 people from 10 countries over the past two decades. The plot project is part the Center for Tropical Forest Science, a global network of large-scale demographic tree plots.

