## Supplementary Information for "Universal abundance fluctuations across microbial communities, tropical forests, and urban populations"

September 13, 2023

**S5 Alternative goodness of fit tests** 7

**S6 Robustness to changing time interval of sampling** 7

**S7 Are forest clusters significantly different?** 8

**S8 Fitting absolute abundance data** 9

**S9 Supplementary figures** 9

#### S1 Solving the Stochastic Linear-Response Model

The stochastic Linear-Response model is given by

$$\frac{dx_i}{dt} = \frac{x_i^*}{\tau_i} - \frac{x_i}{\tau_i} + \sqrt{2\sigma_i x_i}\,\eta_i(t), \tag{S1}$$

where $x_i^*$ is the equilibrium abundance, $\tau_i$ sets timescale of the deterministic forces, $\sigma_i$ determines the strength of population fluctuations, and $\eta(t)$ is delta-correlated Gaussian noise or white noise ($\langle\eta(t)\eta(t\prime)\rangle = \delta(t - t\prime)$).

The Fokker-Planck equation for the probability density function corresponding to the above stochastic differential equation, by the Ito prescription,

$$\frac{\partial P_i(x,t)}{\partial t} = -\frac{\partial}{\partial x}\left[\left(\frac{x_i^* - x}{\tau_i}\right) P_i(x,t)\right] + \sigma_i \frac{\partial^2}{\partial x^2}\left[x P_i(x,t)\right] \tag{S2}$$

##### S1.1 Steady-state solution

At steady-state, the time derivative is zero, and so we get an equation for the steady-state probability density function $P_{\mathrm{ss},i}(x)$ :

$$\left(\frac{x_i^* - x}{\tau_i}\right) P_{\mathrm{ss},i}(x) = \sigma_i \frac{\partial}{\partial x}\left[x P_{\mathrm{ss},i}(x)\right]. \tag{S3}$$

To solve this equation, we define $Q_i(x) = x P_{\mathrm{ss},i}(x)$ and simplify to get

$$\frac{1}{Q_i(x)}\frac{\partial Q_i(x)}{\partial x} = \frac{1}{\sigma_i \tau_i}\left(\frac{x_i^*}{x} - 1\right). \tag{S4}$$

This equation can be integrated to get

$$Q_i(x) = c x^{\frac{x_i^*}{\sigma_i \tau_i}} e^{-\frac{x}{\sigma_i \tau_i}}, \tag{S5}$$

where $c$ is an undetermined constant of integration. We can now calculate the steady-state pdf using $P_{\mathrm{ss},i}(x) = x^{-1} Q_i(x)$ and calculate the constant of integration from the normalization condition $\int_{-\infty}^{\infty} P_{\mathrm{ss},i}(x)dx = 1$. This gives us the result

$$P_{\mathrm{ss},i}(x) = \frac{(\sigma_i \tau_i)^{-x_i^*/\sigma_i \tau_i}}{\Gamma(x_i^*/\sigma_i \tau_i)} x^{\frac{x_i^*}{\sigma_i \tau_i} - 1} e^{-\frac{x}{\sigma_i \tau_i}}, \tag{S6}$$

which is the Gamma distribution. The properties of the Gamma distribution are well-known. The mean of the Gamma distribution is $x_i^*$ and the ratio of variance to mean-squared is the ENR $\frac{x_i^*}{\sigma_i \tau_i}$.

##### S1.2 Time-dependent solution

We can also obtain the full time-dependent solution of the SLRM. Analytical solutions of this model have been obtained in the context of diffusion processes, bond interest rates, and birth-death models [1, 2, 3]. To obtain the solution, we consider the backward Kolmogorov equation describing the dynamics

$$-\frac{\partial P_i(x,t)}{\partial t} = \left(\frac{x_i^* - x}{\tau_i}\right)\frac{\partial}{\partial x} P_i(x,t) + \sigma_i x \frac{\partial^2}{\partial x^2} P_i(x,t) \tag{S7}$$

We can non-dimensionalize the parameters by changing variable to $z = \frac{x}{\sigma_i \tau_i}$, $\theta = \frac{t}{\tau_i}$ to get

$$-\frac{\partial P_i(z,\theta)}{\partial \theta} = \left(\frac{x_i^*}{\sigma_i \tau_i} - z\right)\frac{\partial}{\partial z} P_i(z,\theta) + z\frac{\partial^2}{\partial z^2} P_i(z,\theta) \tag{S8}$$

We now use the separation of variables ansatz to look for solutions of the form $P(z,\theta) = q_\lambda(z) r_\lambda(\theta)$. Plugging this in, we get

$$\frac{-1}{r_\lambda(\theta)}\frac{\partial r_\lambda(\theta)}{\partial \theta} = \frac{1}{q_\lambda(z)}\left[\left(\frac{x_i^*}{\sigma_i \tau_i} - z\right)\frac{\partial q_\lambda(z)}{\partial z} + z\frac{\partial^2 q_\lambda(z)}{\partial z^2}\right] = -\lambda. \tag{S9}$$

Hence we have $r_\lambda(\theta) = e^{\lambda\theta}$ and an equation for $q_\lambda(z)$:

$$\left(\frac{x_i^*}{\sigma_i\tau_i} - z\right)\frac{\partial q_\lambda(z)}{\partial z} + z\frac{\partial^2 q_\lambda(z)}{\partial z^2} + \lambda q_\lambda(z) = 0. \tag{S10}$$

This is the confluent hypergeometric equation [4]. It has two linearly independent solutions when $\frac{x_i^*}{\sigma_i\tau_i}$ is positive, denoted as $\psi_1(z)$ and $\psi_2(z)$. Hence the general solution is of the form $q_\lambda(z) = c_1\psi_1(z) + c_2\psi_2(z)$.

The two functions are defined as $\psi_1(z) = U(-\lambda, \frac{x_i^*}{\sigma_i\tau_i}; z)$ and $\psi_2(z) = e^z U(\lambda + \frac{x_i^*}{\sigma_i\tau_i}, \frac{x_i^*}{\sigma_i\tau_i}; z)$, where $U(a, b; z)$ is defined by the integral $U(a, b; z) = \frac{1}{\Gamma(a)}\int_0^\infty e^{-zs}s^{a-1}(1+a)^{b-a-1}ds$.

Imposing finite moments for the solution sets $c_2 = 0$ and $\lambda$ to take integer values. At these integer values, the eigen functions are Laguerre polynomials. Following Ref. [3], we can solve for the coefficients assuming a reflecting boundary condition at $z = 0$ and delta function initial condition at $t = 0$, $\delta(x - x_0)$.

The time-dependent solution thus reads,

$$P_i(x, t|x_0, 0) = \frac{1}{\sigma_i\tau_i\left(1 - e^{-t/\tau_i}\right)}\left(\frac{x}{x_0 e^{-t/\tau_i}}\right)^{-\frac{1}{2}+\frac{x_i^*}{2\sigma_i\tau_i}} \exp\left[-\frac{x + x_0 e^{-t/\tau_i}}{\sigma_i\tau_i\left(1 - e^{-t/\tau_i}\right)}\right] I_{\frac{x_i^*}{\sigma_i\tau_i}-1}\left[\frac{2\sqrt{x x_0 e^{-t/\tau_i}}}{\sigma_i\tau_i\left(1 - e^{-t/\tau_i}\right)}\right]. \tag{S11}$$

This is, in fact, the non-central chi-squared distribution, $\chi^2\left(\frac{2x}{\sigma_i\tau_i\left(1-e^{-t/\tau_i}\right)}; \frac{2x_i^*}{\sigma_i\tau_i}, \frac{2x_0 e^{-t/\tau_i}}{\sigma_i\tau_i\left(1-e^{-t/\tau_i}\right)}\right)$.

#### S1.3 The distributions of fold-change and logarithmic fold-change

We can use the time-dependent solution (Eq. (S11)) and the steady-state distribution ((Eq. (S6)) to calculate the probability of abundance fluctuations. Specifically, we will first calculate the probability distribution of the fold-change in a time interval $\Delta t$, $\rho = x(\Delta t)/x(0)$.

The fold-change distribution, $P_{\mathrm{fc},i}(\rho, \Delta t)$, is given by

$$P_{\mathrm{fluc},i}(\rho, \Delta t) = \int_0^\infty dx(0)\int_0^\infty dx(\Delta t) P_i(x(\Delta t), t + \Delta t|x(0), 0) P_{ss}(x(0))\delta\left(\rho - \frac{x(\Delta t)}{x(0)}\right). \tag{S12}$$

Plugging the expressions from Eqs. (S11) and (S6) in the above, the integral is evaluated as in Ref. [3] to get

$$P_{\mathrm{fc},i}(\rho, \Delta t) = \frac{2^{\frac{x_i^*}{\sigma_i\tau_i}-1}}{\sqrt{\pi}}\frac{\Gamma\left(\frac{x_i^*}{\sigma_i\tau_i}+\frac{1}{2}\right)}{\Gamma\left(\frac{x_i^*}{\sigma_i\tau_i}\right)}\frac{(\rho+1)}{\rho}\frac{\left(e^{\Delta t/\tau_i}\right)^{\frac{x_i^*}{2\sigma_i\tau_i}}}{1 - e^{-\Delta t/\tau_i}} \left(\frac{\sinh\left(\frac{\Delta t}{2\tau_i}\right)}{\rho}\right)^{\frac{x_i^*}{\sigma_i\tau_i}+1}\left(\frac{4\rho^2}{(\rho+1)^2 e^{\Delta t/\tau_i} - 4\rho}\right)^{\frac{x_i^*}{\sigma_i\tau_i}+\frac{1}{2}} \tag{S13}$$

We can now change variables from the fold-change $\rho = \frac{x(\Delta t)}{x(0)}$ to logarithmic fold-change $\lambda = \ln\frac{x(\Delta t)}{x(0)}$ by using $P_{\mathrm{fluc},i}(\lambda, \Delta t) = e^\lambda P_{\mathrm{fc},i}(e^\lambda, \Delta t)$. Simplifying, we get the expression for the Logarithmic Fold-change distribution (LFD),

$$P_{\text{fluc},i}(\lambda,\Delta t) = \frac{\Gamma\left(\frac{x_i^*}{\sigma_i\tau_i}+\frac{1}{2}\right)}{2\sqrt{\pi}\Gamma\left(\frac{x_i^*}{\sigma_i\tau_i}\right)}\left(e^{\frac{\Delta t}{\tau_i}}-1\right)^{\frac{x_i^*}{\sigma_i\tau_i}} e^{\frac{\Delta t}{2\tau_i}}$$

$$\cosh\frac{\lambda}{2}\left(\frac{1}{e^{\frac{\Delta t}{\tau_i}}(\cosh\frac{\lambda}{2})^2-1}\right)^{\frac{x_i^*}{\sigma_i\tau_i}+\frac{1}{2}}. \quad \text{(S14)}$$

The corresponding cumulative distribution function can be computed as well, to make sampling the distribution easier. The cumulative distribution function corresponding to $P_{\text{fluc},i}(\lambda,\Delta t)$, $CDF_{\text{fluc},i}(\lambda,\Delta t)$ is given by

$$CDF_{\text{fluc},i}(\lambda,\Delta t) = \frac{1}{2}+\frac{\Gamma\left(\frac{x_i^*}{\sigma_i\tau_i}+\frac{1}{2}\right)}{\sqrt{\pi}\Gamma\left(\frac{x_i^*}{\sigma_i\tau_i}\right)}\sinh\frac{\lambda}{2}\left(1-e^{-\frac{\Delta t}{\tau_i}}\right)^{-\frac{1}{2}} {}_2\mathrm{F}_1\left[\frac{1}{2};\frac{x_i^*}{\sigma_i\tau_i}+\frac{1}{2};\frac{3}{2};\frac{-\sinh^2\frac{\lambda}{2}}{1-e^{-\frac{\Delta t}{\tau_i}}}\right] \quad \text{(S15)}$$

where ${}_2\mathrm{F}_1$ denotes the hypergeometric function.

### S2 The SLRM as a linearization of a nonlinear model

The most straightforward interpretation of the SLRM is that it arises from a more complicated nonlinear model as a linearization around the model equilibrium. Many ecological population models converge to a unique stable equilibrium (see Table 1 in [5]). If we assume that the deterministic part of the dynamics is described by an equation $\frac{dx_i}{dt} = f(\vec{x})$, where $\vec{x}$ is the vector of species abundances in the system and has an equilibrium at $\vec{x} = \vec{x}^*$ . Then deviations from the equilibrium gives rise to a linear restoring force of the form $\nabla f(\vec{x}^*)\cdot(\vec{x}^*-\vec{x})$, where $\nabla$ refers to the gradient operator. The SLRM is obtained when we assume that the restoring force to equilibrium on species $i$ is dominated by its own deviation from equilibrium i.e., we neglect contributions from coupling between species. Neglecting interactions allows us to restrict an otherwise large number of model parameters.

This does suggest a natural extension of the SLRM that accounts for species interactions, as discussed in the main text. A simple way to incorporate species interactions would be to augment the deterministic part of the SLRM with contributions from the other species $\nabla f(\vec{x}^*)\cdot(\vec{x}^*-\vec{x})$. This would result in an extension of the SLRM described by

$$\frac{dx_i}{dt} = \nabla f(\vec{x}^*)\cdot(\vec{x}^*-\vec{x}) + \sqrt{2\sigma_i x_i}\,\eta_i(t). \quad \text{(S16)}$$

Investigating this model and applying it to data is an interesting direction for future work.

#### S2.1 Example of SLRM arising from a model with interactions

Here we show an example scenario of how the SLRM might arise from a population dynamics model with interactions without linearizing the dynamics. Specifically, we consider a generalized Lotka-Volterra (gLV) model with immigration and demographic noise:

$$\frac{dx_i}{dt} = m_i + g_i x_i - \sum_j A_{ij}x_i x_j + \sqrt{2\sigma_i x_i}\,\eta(t), \quad \text{(S17)}$$

where $m_i$ is the immigration, $g_i$ is the growth rate, the matrix $A$ measures inter-species interactions, $\sigma_i$ measures strength of demographic noise, and $\eta(t)$ is delta-correlated Gaussian noise.

The limit we consider, reminiscent of a mean-field approximation in physics, is obtained by replacing the inter-species interaction $A_{ij}$ with an average interaction strength $B_i = \frac{1}{S}\sum_j A_{ij}$, where $S$ is the number of species. This leads to a low-rank approximation of the interaction matrix, with model dynamics being described by

$$\frac{dx_i}{dt} = m_i + g_i x_i - B_i x_i \sum_j x_j + \sqrt{2\sigma_i x_i}\,\eta(t). \tag{S18}$$

For relative abundances, $\sum_j x_j = 1$ and neglect noise correlations in the high diversity limit (alternatively for absolute abundances, we neglect fluctuations in the total population size so that $\sum_j x_j = c$ is a constant). With this, we get

$$\frac{dx_i}{dt} = m_i + g_i x_i - B_i x_i c + \sqrt{2\sigma_i x_i}\,\eta(t), \tag{S19}$$

which can easily be recast into form of the SLRM (Eq. (S1)) by re-expressing the parameters as $m_i = \frac{x_i^*}{\tau_i}$ and $g_i - B_i c = -\frac{1}{\tau_i}$. Thus, we obtain the SLRM from the model with interactions in a special limiting scenario. Note that this is intended to serve only as an example scenario where SLRM is derived from an interacting model without linearization. Whether the SLRM may arise as an effective model of more complex interacting models in other scenarios is an interesting topic for future work.

#### S3 The SLRM for relative vs absolute abundances.

The SLRM equations in the main text were written in terms of the relative abundances $x_i$, which is also the data we analyze. Many common population models, however, are written in terms of the absolute abundance $n_i$ instead. Here we show that when fluctuations in the total population size can be neglected, the SLRM for relative abundances implies an SLRM for absolute abundances as well.

We start by assuming the absolute abundances of a sector/species, $i$, in a city/community $c$ is described by an SLRM. Specifically, the dynamics is given by

$$\frac{dn_{ci}}{dt} = \frac{n_{ci}^*}{\tau_{ci}} - \frac{n_{ci}}{\tau_{ci}} + \sqrt{2\sigma_{ci}^{(n)} n_{ci}}\,\eta_{ci}(t), \tag{S20}$$

where without loss of generality we have assumed that the demographic parameters depend on both $c$ and $i$. The equation for the relative abundances $x_{ci} = n_{ci}/N_c$, where $N_c$ is the total population of the city/community is then given by

$$\frac{dx_{ci}}{dt} = \frac{x_{ci}^*}{\tau_{ci}} - \frac{x_{ci}}{\tau_{ci}} + \sqrt{2x_{ci}\sigma_{ci}^{(n)}/N_c}\,\eta_{ci}(t), \tag{S21}$$

if we neglect variation in the total population $\frac{dN_c}{dt}$. This variation in total abundances could be small due to some unmodelled global feedback that leads to the approximated cancellation of positive and negative fluctuations of many population categories. We can now redefine the noise term by using $\sigma_{ci}^{(x)} = \sigma_{ci}^{(n)}/N_c$. With this redefinition, the equation becomes

$$\frac{dx_{ci}}{dt} = \frac{x_{ci}^*}{\tau_{ci}} - \frac{x_{ci}}{\tau_{ci}} + \sqrt{2x_{ci}\sigma_{ci}^{(x)}}\,\eta_{ci}(t), \tag{S22}$$

which is identical to Eq.(S20), except with $\sigma_{ci}^{(x)}$. Thus an SLRM for relative abundances is equivalent to an SLRM for absolute abundances when the fluctuations in the total population size can be neglected. In general, these fluctuation may not be neglected and transforming the SLRM to absolute abundances would need to assume a dynamical equation for $\frac{dN_c}{dt}$.

In the main text, we have made an additional simplifying assumption. When analyzing city data, we assumed that the parameters $\tau_{ci}$,$x_{ci}^*$, and $\sigma_{ci}^{(x)}$ depend only on the sector $i$ and are independent of the city $c$. Analogous assumptions were applied to forest data. For microbiome analysis, we consider longitudinal data from only a single community $c$ and do not integrate data from different communities; hence such an assumption was not required. Also, note that assuming $\sigma_{ci}^{(x)}$, depend only on the sector $i$ implies that $\sigma_{ci}^{(n)}/N_c$ is a constant independent of city size. This makes fluctuations important for big and small cities alike.

### S4 Literature review

In this section, we summarize works examining the abundances and temporal fluctuations of the three complex populations we study.

#### S4.1 Microbiome

In the context of the microbiome time-series data, Refs. [6, 7, 8] have proposed and applied the Stochastic Logistic Model (SLM) to microbiome data. The Stochastic Logistic Model is a three-parameter model that combines a nonlinear, deterministic part describing logistic growth with environmental stochasticity. In Fig.7 in the main text, we compare the performance of the Stochastic Linear Response Model (SLRM) and the Stochastic Logistic Model (SLM) in explaining the statistical distributions of abundances and its fluctuations in all three systems. We found that the SLRM outperformed the SLM the majority of cases (17 out of the 18 employment sectors, 72 out of 85 microbial species, and 2 out of the 4 forest clusters).

A number of studies have also fit microbiome data with more complex models such as Lotka-Volterra and consumer-resource models [9, 10]. Ref. [11] examined the Logarithmic Fold-change Distribution (LFD) in microbiome data, aggregated over all species. They fit the LFD with a Laplace distribution and used this to narrow the parameter space of consumer-resource models they were investigating. In Fig. S2, we compare the fits of the empirical LFD in all three systems Laplace and Gaussian candidate distributions. The SLRM outperformed the other distributions in majority of cases. Finally, Ref. [12] proposed a birth-death-migration model for microbiomes but do not apply it to data.

#### S4.2 Forests

Using tropical forest data to investigate population fluctuations was catalyzed by two key innovations: empirically, the development of whole-plot censuses over multiple years by the CTFS [13] provided data that can be used to analyze population fluctuations at scale. In the area of theoretical modeling, there was the development of the neutral theory of biodiversity [14, 15, 16], and its potential explanation of static patterns of diversity like the species abundance distribution and spatial turnover [17, 18, 19, 20]. This naturally led to the question of whether the combination of competition for space and demographic noise could explain temporal patterns alongside spatial patterns. Azaele et al [3] used the SLRM formulation to test whether square-root (i.e. demographic-like) noise could potentially explain patterns of fluctuations in the Barro Colorado Island plot, treating all species as governed by the same parameter values. Our model follows a similar approach, though we allow for populations to take on different parameter values according to which of four height niches they belong to.

Subsequent developments in the analysis of forest population fluctuations [21, 22, 23] considered the possibility that environmental stochasticity [24, 25], may provide a more comprehensive explanation. These models introduce independent, uncorrelated environmental noise for each species alongside demographic noise and competition for space, providing a better fit to several data sets than demographic noise alone, though at the conceptual expense of introducing unidentified environmental variables. We also note that the metrics used to assess the fit to fluctuation sizes differ from those used here and by Azaele et al, and here in our analysis, and in our own analysis we show that square-root noise alone may provide an adequate explanation of tropical forest fluctuations.

##### S4.3 Cities

For the dynamics of sectoral employment in cities, work has been mostly limited to regional effects, sector-specific analyses, or understanding the effect of total city size on employment composition by sector in the city [26, 27, 28, 29]. In a broader context, simple mathematical models have been successfully applied to explain the distribution of population sizes across cities and population migration patterns [30, 31, 32].

#### S5 Alternative goodness of fit tests

In addition to the likelihood based goodness fit test presented in Fig.4, we applied two alternative goodness of fit tests to the data: a test based on the Jensen-Shannon Distance (JSD) [33, 34] and the Kolomogorov-Smirnov (KS) test [34, 35].

The Jensen-Shannon Distance (JSD) measures the distances between two probability distributions. The JSD-based test was performed similarly to the likelihood based test. 1000 Random samples of the same size of the fitted distribution were generated at the estimated parameters. Then the JSD between the observed data and theoretical distribution was compared to JSDs of the random samples and the theoretical distribution. We passed the fit if the JSD between observed data and theory was was smaller than at least 1% of the simulated data (i.e., rejected at a 1% threshold). 13 of 18 sectors, 43 of 85 microbial species, and 3 of 4 forest clusters passed the JSD test. Visualizing the abundance distributions of species that were rejected by the test, we found that rejection was sometimes driven by just one or two outlier data points where the species was recorded as being highly abundant (see Fig. S7C,G). Whether this signifies the ground truth or measurement error is an open question.

The one-sample KS test is a non-parametric test that compares the observed data with a reference distribution (theoretical prediction). It generates a p-value associated with each comparison. 15 of 18 sectors, 55 of 85 microbial species, and 3 of 4 forest clusters passed the KS test at a 1% threshold.

#### S6 Robustness to changing time interval of sampling

To test the robustness of our results, we investigated the effect of changing the time interval between consecutive data points over which the Log Fold-change was calculated, $\Delta t$. This could be performed for the microbiome and cities because there were enough time points in both these data sets to allow us to vary $\Delta t$. To implement the test, we first obtained the empirical LFD for different values of sampling interval $\Delta t$. The theoretical LFD, $P_{\mathrm{fluc}}$, depends on two parameters, the ENR and $\Delta t/\tau$. We obtained the ENR from fitting the empirical abundance distribution (SSD) and then fit the empirical LFD to obtain $\Delta t/\tau$ as $\Delta t$ was varied.

Fig. S8 shows the results of this analysis for employment data in cities. We found that the inferred parameter combination $\Delta t/\tau$ increased approximately linearly with increasing $\Delta t$ as we expect (Fig. S8A). In other words, the inferred parameter $\tau$ is approximately independent of the sampling interval $\Delta t$ indicating a robust fit (Fig. S8B).

In contrast to the employment data, analysis of the microbiome data showed that the inferred parameter combination $\Delta t/\tau$ increased slower than linearly with $\Delta t$ (Fig. S9A). Furthermore, at large $\Delta t$ the inferred parameters of a few species oscillated. This spurious behavior can be understood by examining how the predicted LFD $P_{\mathrm{fluc}}$ changes as we vary $\Delta t/\tau$ (Fig. S9B). The distribution does not change much once $\Delta t$ approaches $\tau$ or $\frac{\Delta t}{\tau} \gtrapprox 1$. Intuitively, this happens because the system equilibrates over a time scale of $\tau$. The fold-change distribution thus remains approximately unchanged for timescales greater than $\tau$. Since the inferred equilibration timescale for microbes is not significantly larger than the sampling interval, this makes the parameter estimation difficult as we increase $\Delta t$. Fig. S9C-F shows the data corresponding to one of the microbes that exhibited a spurious oscillation in its inferred parameter. Small changes in the data led to large changes in the inferred parameter because $\frac{\Delta t}{\tau} \gtrapprox 1$.

#### S7 Are forest clusters significantly different?

In our analysis of BCI data, we classified species into four distinct clusters based on their maximum height. This was done under the assumption that species with similar height have similar access to light, in terms of level, variability, or horizontal uniformity, and hence are likely to be described the similar parameters [36]. We utilized the results from the quantitative analysis of BCI data in Ref. [37] to assign species to four distinct height clusters: shrubs, understory treelets, midstory trees, and canopy trees.

In this section, we examine how different the extent of dissimilarity in the inferred parameters and emergent behavior of the four different height clusters. For this, we conducted goodness of fit tests to see if the parameters of individual tree clusters are differed significantly from each other. Specifically, for each pair of clusters, eg. ($c_1$=shrubs, $c_2$=canopy), we took the fit parameters of cluster $c_1$ and computed the likelihood and JSD metrics using the data from cluster $c_2$. We compared this to the likelihood and JSD metrics obtained from 100 random samples from the LFD and 1000 samples from the SSD. Note that ($c_1$=shrubs, $c_2$=canopy) is different from ($c_1$=canopy, $c_2$=shrubs) and so there are 12 such pairs we can test.

At a 5% threshold for the LFD, the likelihood and JSD tests rejected 3 and 6 pairs respectively. All of the rejected pairs involved comparing a shrub with one of the other three clusters. Fig. S13 A indicates that the primary difference us that shrubs have a significantly shorter equilibration timescale than the other clusters. This manifest in the observed Log Fold-change data as shrubs experiencing much larger fluctuations than the other three clusters (Fig. S13 B). This might be reflective of the shorter generation times of shrubs. From the fitted distributions shown, we also observe that the fitted parameters of midstory trees and canopy trees are highly similar.

The distinction between shrubs and the other three clusters appeared in the test results of the SSD as well. At a 5% threshold for the SSD, the likelihood and JSD tests rejected 3 and 6 pairs respectively. Shrubs were involved in 6 of the 9 rejected pairs. Thus, in terms of the emergent properties analyzed in this study, shrubs form the most distinct height cluster among the BCI forest species.

#### S8 Fitting absolute abundance data

In our analyses so far, we used relative abundance data in all three systems. This was done for a few reasons. First, sequencing does not provide absolute abundance data in the microbiome data, and so this allows us to treat data from all three systems identically. Second, our definition of relative abundances in employment, as the employment in a sector in a city divided by the total employment in the city, allows us to ignore the large variation in the absolute sizes of cities [30]. Third, it allows us to ignore the effects of changing overall population sizes (Fig. S10A,B).

Despite the trends in overall population sizes (Fig. S10A,B), the fluctuations in the logarithmic fold-change of species/sectors calculated as $\frac{n_i(t+\Delta t)}{n_i(t)}$ are still large compared to the mean (Fig. S10C,D). Therefore, we tried fitting the empirical LFD calculated from absolute abundance data of species in forests and sectoral employment in cities (Fig. S11). The SLRM was still able to fit the data well. It outperformed other candidate distributions in all employment sectors in the city data and 2 of the 4 forest clusters, as compared to outperforming candidate distributions in all employment sectors and all 4 forest clusters when using LFD from relative abundance data. This slight decrease in performance against candidate distributions for the forest data may indeed be due to the temporal drift of overall population sizes.

We also tried to fit the abundance distribution to the SSD predicted by the SLRM for forest data. The SLRM was able to fit the data reasonably well (Fig. S12). We did not fit the abundance distribution of absolute sectoral employment across cities because we expect the variation in population sizes across cities to play a dominant role in this analysis. City sizes are known to vary widely and display a power-law like behavior [30]. Indeed, this was one of our motivations behind normalizing the sectoral employment in a city by the total employment in the same city.

#### S9 Supplementary figures

| NAICS code | Sector name (full) | Sector name (short) |
|---|---|---|
| 11 | Agriculture, Forestry, Fishing and Hunting | Agriculture |
| 21 | Mining, Quarrying, and Oil and Gas Extraction | Mining |
| 22 | Utilities | Utilities |
| 23 | Construction | Construction |
| 31-33 | Manufacturing | Manufacturing |
| 42 | Wholesale Trade | Wholesale |
| 44-45 | Retail trade | Retail trade |
| 48-49 | Transporation | Transporation |
| 51 | Information | Information |
| 52 | Finance and Insurance | Finance |
| 53 | Real Estate and Rental and Leasing | Real Estate |
| 54 | Professional, Scientific, and Technical Services | Professional |
| 55 | Management of Companies and Enterprises | Management |
| 56 | Administrative and Support and Waste Management and Remediation Services | Administrative |
| 61 | Educational Services | Educational |
| 62 | Health Care and Social Assistance | Health |
| 71 | Arts, Entertainment, and Recreation | Arts |
| 72 | Accommodation and Food Services | Accommodation |
| 81 | Other Services (except Public Administration) | Other |
| 92 | Public Administration | Public Administration |
| 99 | Industries not classified | unclassified |

Table S1: **Table of NAICS categories at two digit resolution.** The shortened version of the NAICS category names are used in the text for clarity and brevity.

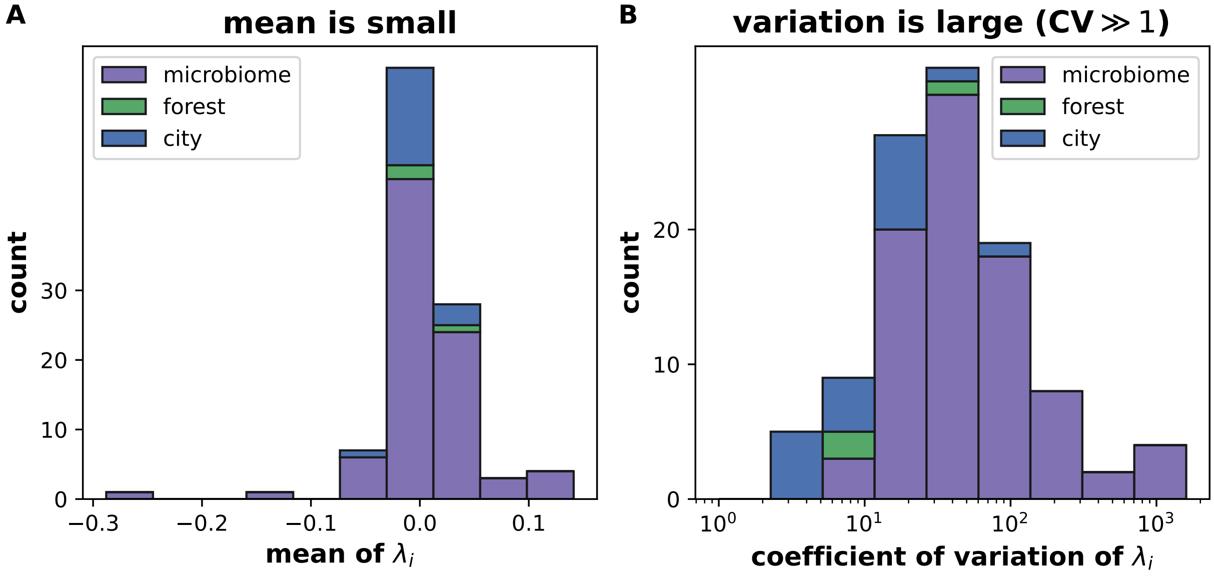

Figure S1: **The mean logarithmic fold-change is much smaller than its standard deviation.** A) The mean logarithmic fold-change $\langle\lambda_i\rangle$ of each trajectory is smaller than the values of $\lambda_i$ shown in Fig.2. B) The coefficient of variation (ratio of standard deviation to mean) is larger than one in all data sets. This prompts us to neglect the small change in the mean in comparison to the large fluctuations between consecutive time points. Both panels show stacked histograms.

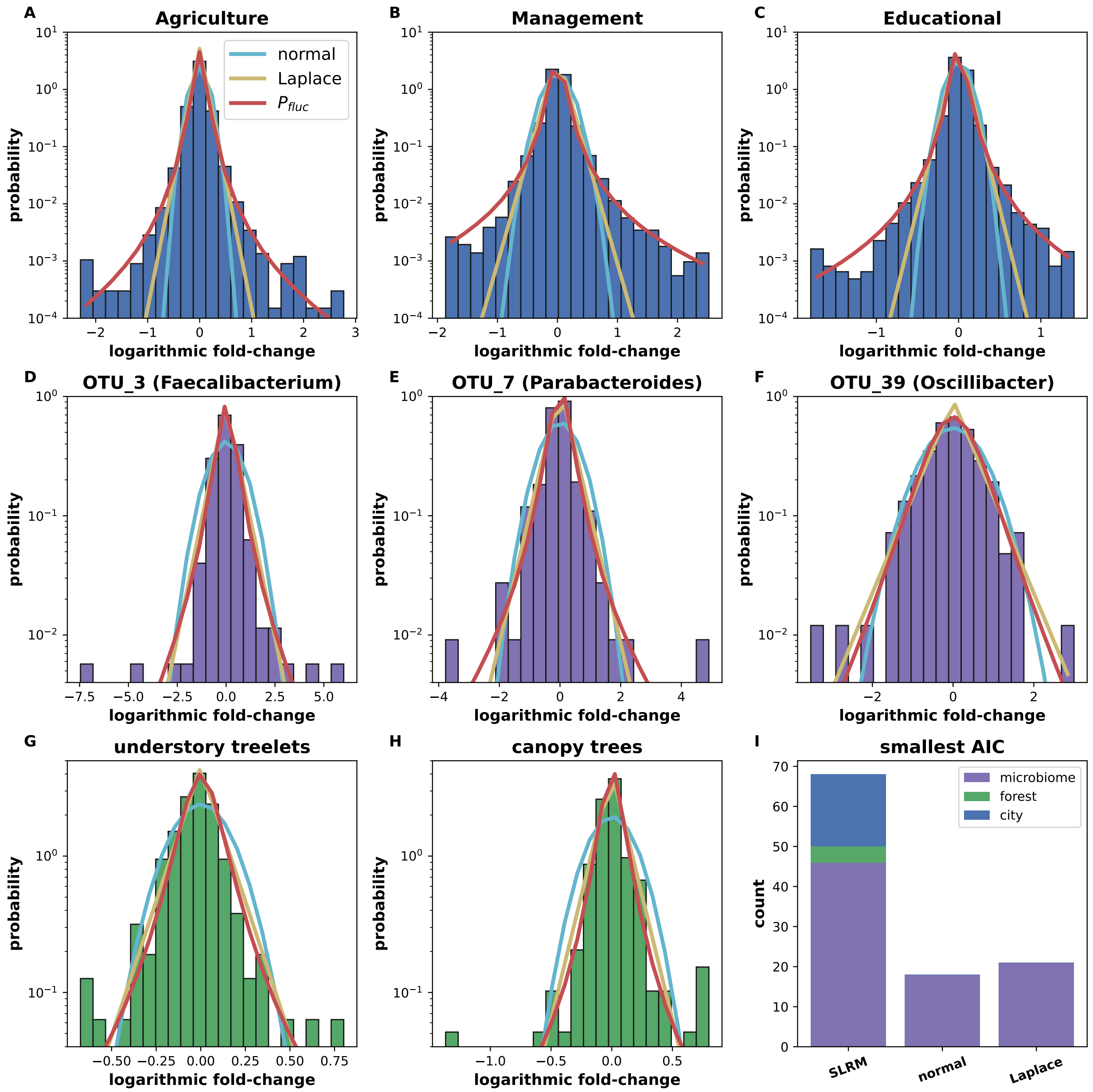

Figure S2: **Model outperforms other candidate distributions in majority of cases.** The empirical Logarithmic Fold-change Distribution (LFD) is shown for three employment sectors **(A-C)**, three microbial species **(D-F)**, and two tree height clusters (**(G,H)**. The LFD is fit by three candidate distributions: normal(cyan), Laplace (yellow), and the prediction from the SLRM, $P_{\mathrm{fluc}}$ (red). **(I)**The SLRM prediction has the smallest Akaike Information Criterion [38] value in the majority of the cases, as show in the stacked histogram. Note that all fitting distributions assumed zero mean, in agreement with our observation of negligible drift (Fig.S1).

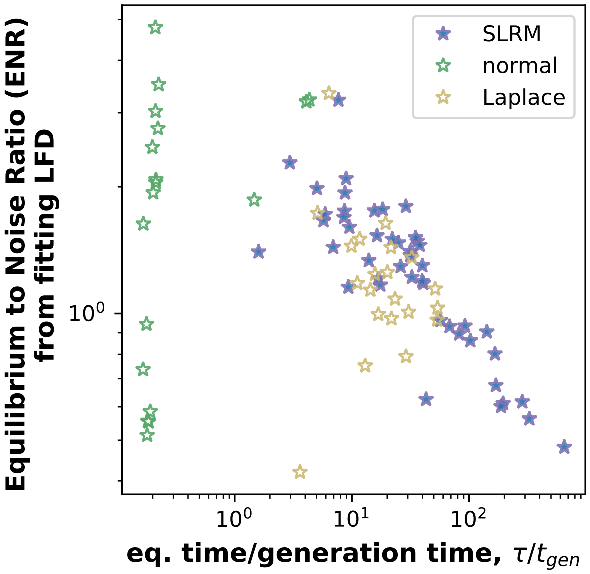

Figure S3: **The microbial species labelled by the model best explaining their logarithmic fold-change distribution.** The SLRM fits the empirical LFD best in the majority of cases (46/85 fits), as measured by the Aikake Information Criterion. Notably, all of the outliers at small $\tau$ values in Fig.3 are better fit by a normal distribution.

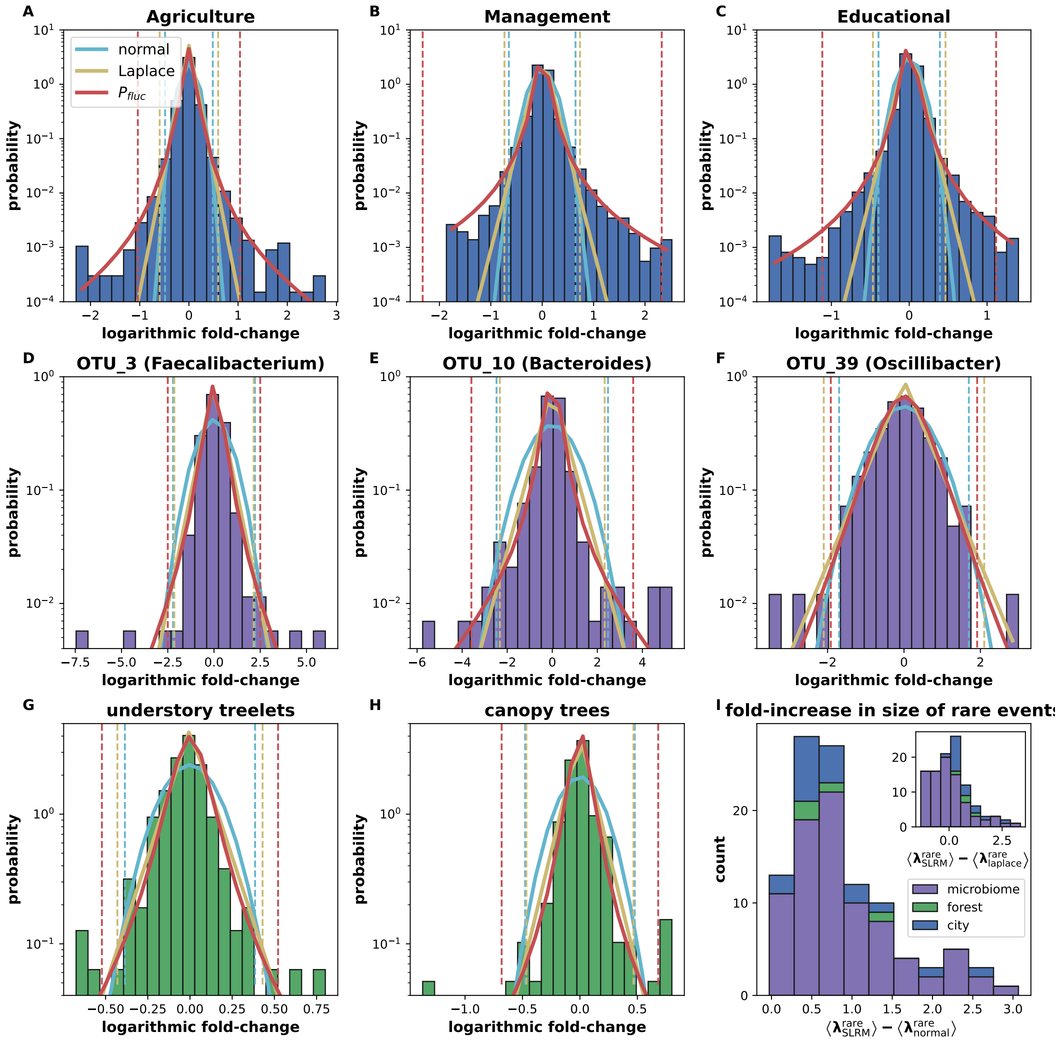

Figure S4: **Difference in size of rare of events estimated by SLRM, Laplace and Normal distribution fits.** **A-H** Examples of Fits of the empirical LFD by the three candidate distributions: SLRM (red), Laplace (cyan), Normal (yellow). Dashed lines bound the central region where majority of the events take place (98% for microbes and forests, 99.8% for cities), as predicted by the three distributions (colored to match). The region outside the dashed lines illustrates the rare, large fluctuations. **I** The error in the estimated logarithmic fold-change of rare, large fluctuations when using alternative distributions instead of SLRM. The size of rare, large fluctuations estimated by each distribution is defined as the expected value of logarithmic fold-change $\lambda$ in the right tail of the distribution (to the right of the dashed lines in panels A-H), $\langle \lambda^{\text{rare}}_{\text{dist.}} \rangle$. The histogram shows the difference between the estimated size of large fluctuations by the SLRM $\langle \lambda^{\text{rare}}_{\text{SLRM}} \rangle$ and normal distribution $\langle \lambda^{\text{rare}}_{\text{normal}} \rangle$. The histogram demonstrates that we will consistently underestimate the size of rare events if we use a normal distribution for risk estimation. The inset shows the difference between the estimated size of large fluctuations by the SLRM $\langle \lambda^{\text{rare}}_{\text{SLRM}} \rangle$ and laplace distribution $\langle \lambda^{\text{rare}}_{\text{normal}} \rangle$. Rare events on the right side of the dashed line had 1% chance of occurring for microbes and forests, and 0.1% for cities. The expectation value of a rare event was defined as $\langle \lambda^{\text{rare}} \rangle = \int_l^\infty \lambda f(\lambda) d\lambda / \int_l^\infty f(\lambda) d\lambda$ for probability distribution function $f(l)$ where the lower limit $l$ was defined as the $\lambda$ value demarcating a rare event (dashed line in panels A-H).

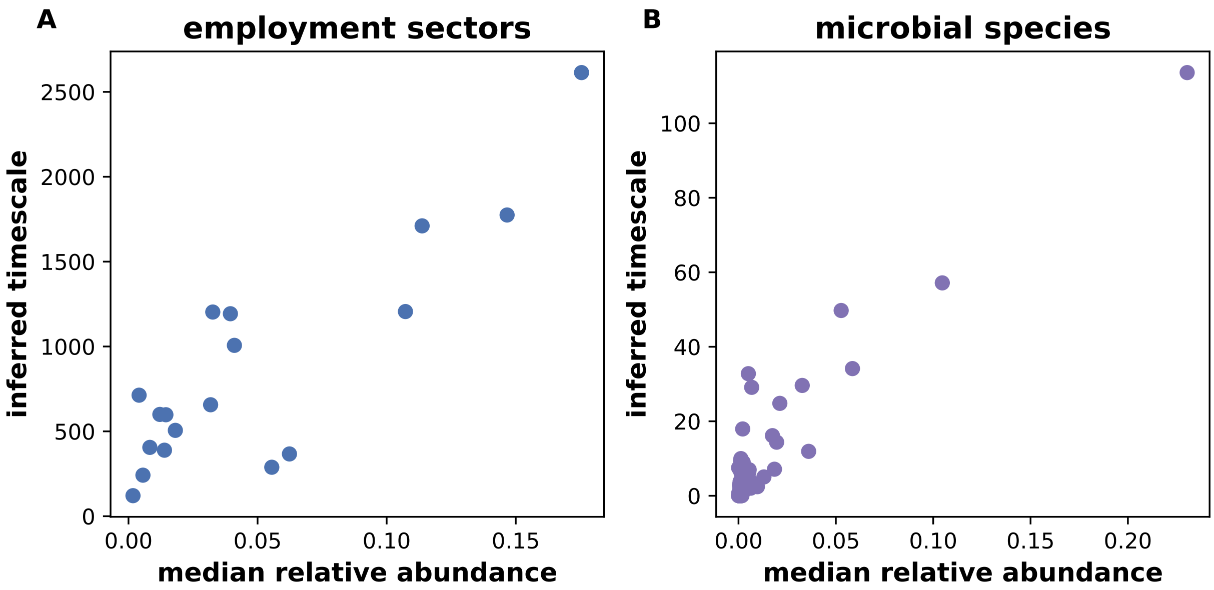

Figure S5: **Inferred timescale, $\tau$ is correlated with abundance of the sector/species.** We found that the inferred timescale from fitting the empirical Log Fold-change Distribution (LFD) is correlated with median abundance of the employment sector **(A)** and microbial species **(B)**. Note that the LFD does not directly contain information about the relative abundance of a species/sector. The median relative employment is calculated across all cities at a given time-point (July 2016) and median relative species abundance is calculated across all time points for the microbiome. Calculated correlations were: pearson's r =0.86, spearman r =0.66 for cities and pearson's r =0.93, spearman r =0.68 for the microbiome. All correlations were statistically significant $p < 0.05$. Axes are plotted on a linear scale because correlations were calculated on a linear scale.

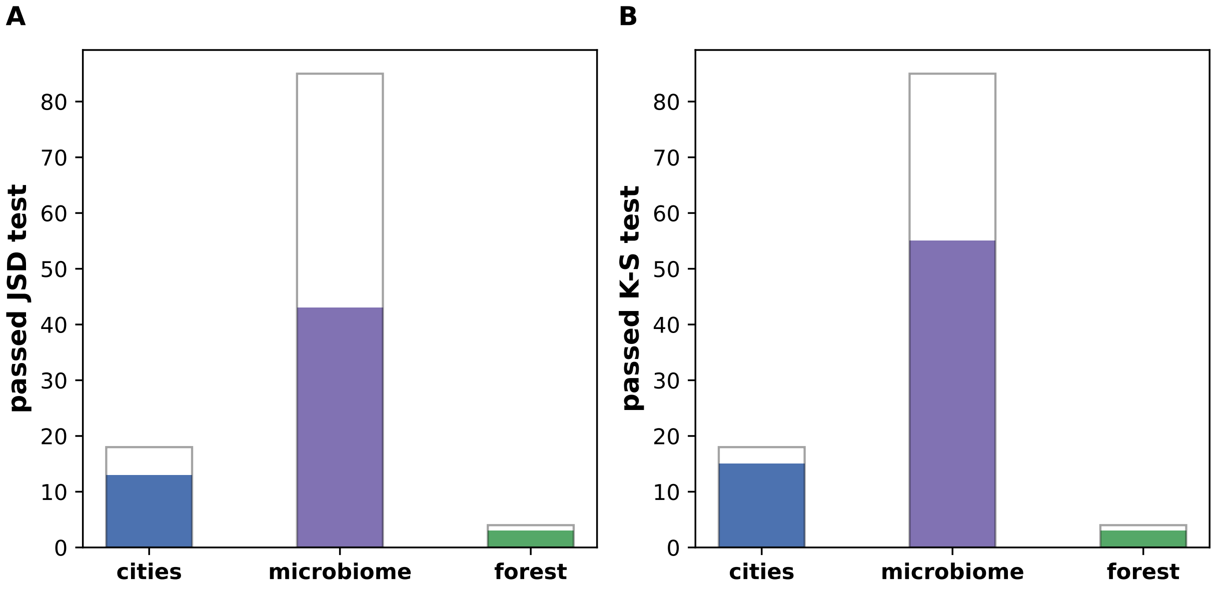

Figure S6: **Alternative goodness of fit tests of the abundance distribution.** In addition to the likelihood based goodness fit test presented in Fig.4, we applied two alternative goodness of fit tests to the data: a test based on the Jensen-Shannon Distance (JSD) [33, 34] and the Kolomogorov-Smirnov (KS) test [34, 35]. A majority of species in each system passed the tests in each system. The species/sectors/clusters that passed the JSD test **(A)** and KS test **(B)** in each system are shown in solid colors. The background shows the total number of species in the systems. In particular, 15 of 18 sectors, 55 of 85 microbial species, and 3 of 4 forest clusters passed the KS test; 13 of 18 sectors, 43 of 85 microbial species, and 3 of 4 forest clusters passed the JSD test.

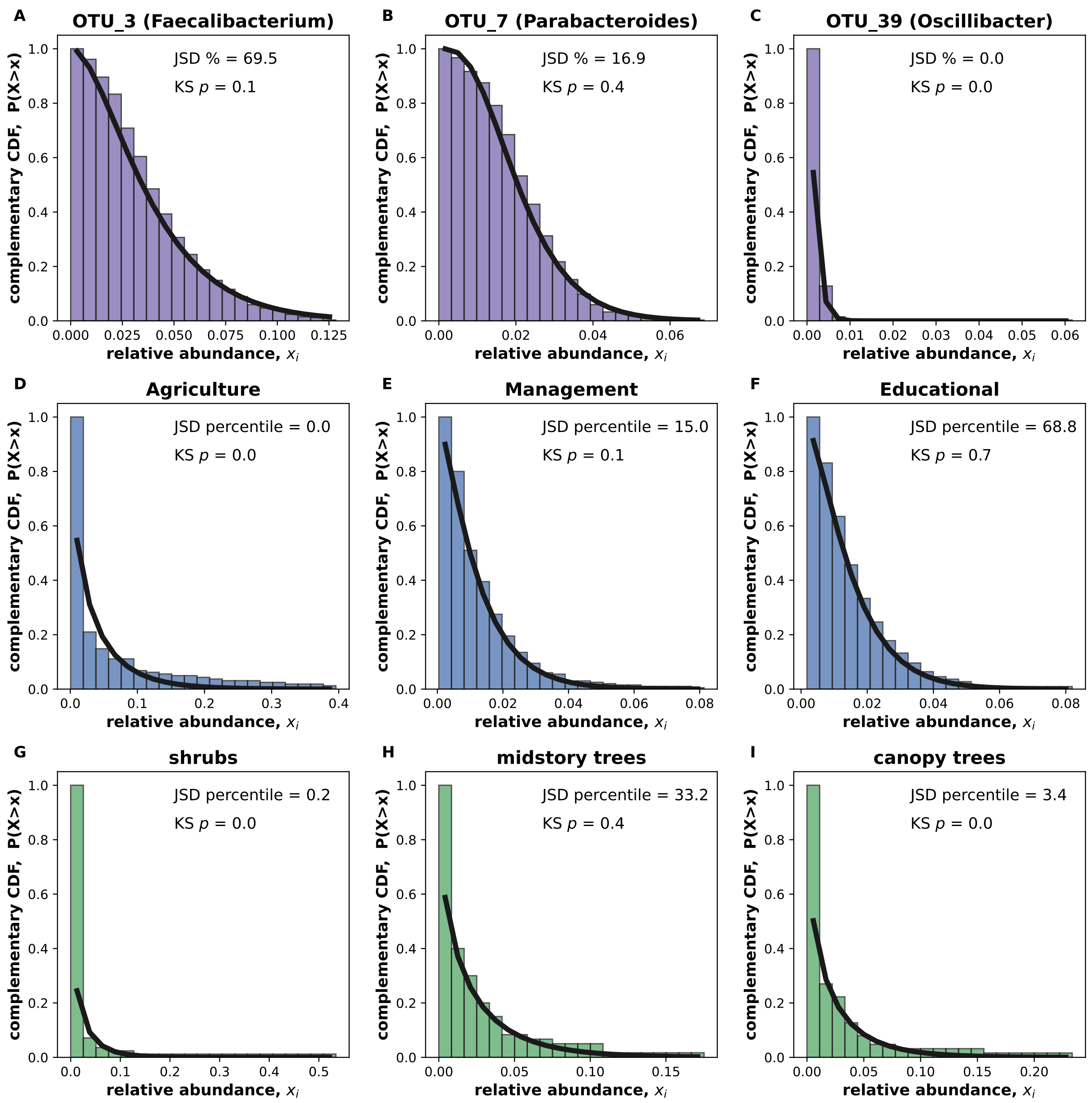

Figure S7: **Abundance distributions with goodness of fit test values. A-I)** The abundance distributions of a few species/sectors/clusters in each system. The percentile score in the JSD-based test and the p-value of the KS test are shown. The fit passed if the percentile score was $> 1\%$ or if $p > 0.01$. Panels C and G illustrate scenarios where the fit was rejected by a single outlier data point near 0.06 and 0.5 respectively.

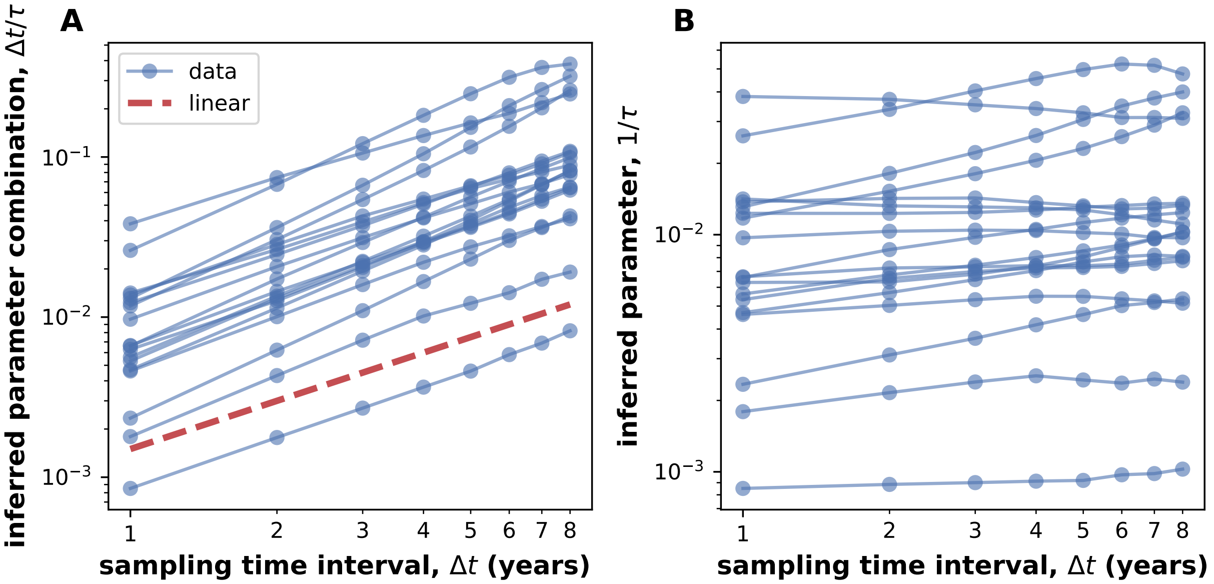

Figure S8: **Scaling of inferred parameters with sampling interval for employment data.** We obtained the empirical LFD at different sampling intervals ($\Delta t$) and fit each one to the theoretical prediction $P_{\text{fluc}}$. **A)** The inferred parameter combination $\Delta t/\tau$ from the fit increased linearly with the sampling interval as expected. **B)**In other words, the inferred timescale, $\tau$, remained approximately constant across fits using data at different $\Delta t$. Sampling intervals $\Delta t$ from 1 to 8 years were used to obtain the empirical LFD. Fits of the LFD were performed at a fixed ENR, which was obtained from fitting the abundance distribution.

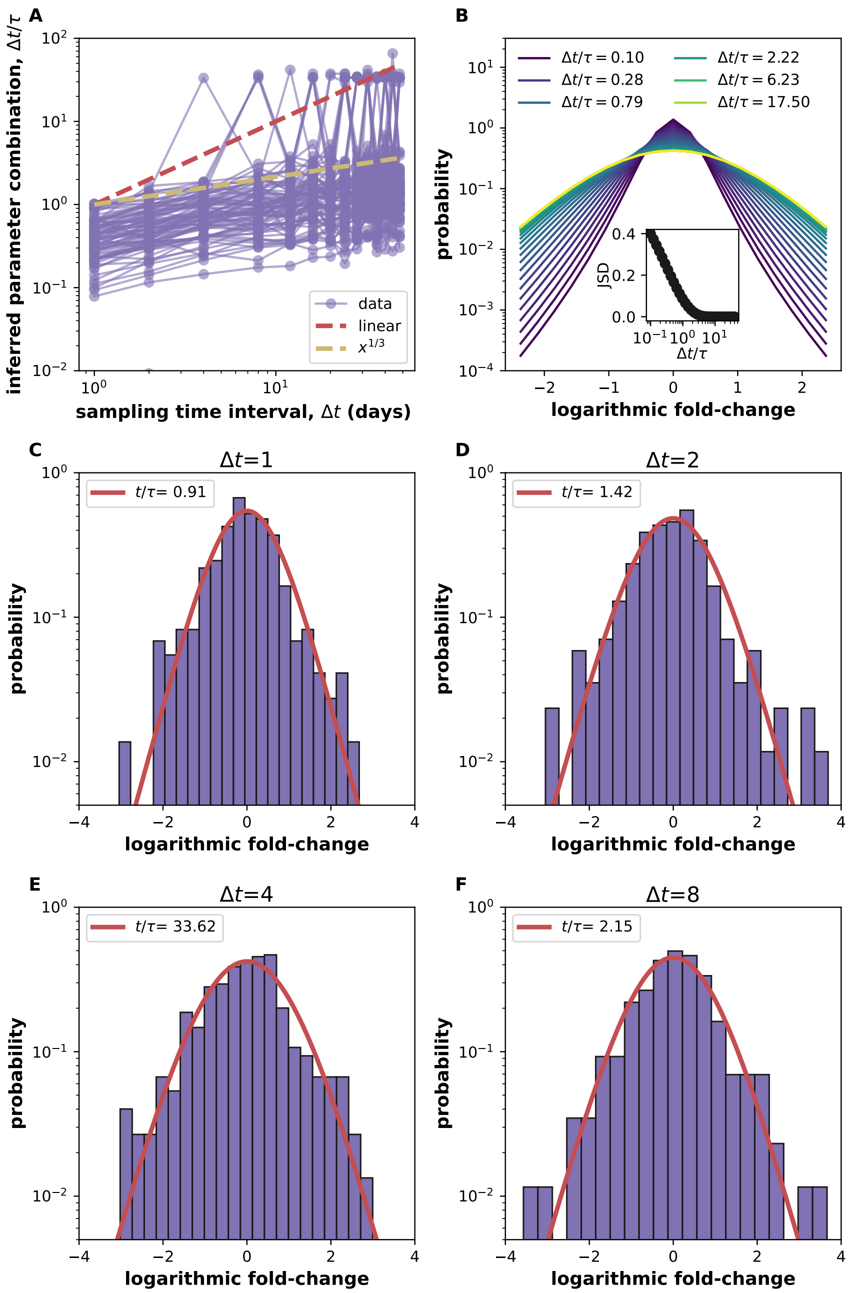

Figure S9: **Scaling of inferred parameters with sampling interval for microbiome data.** We obtained the empirical LFD at different sampling intervals ($\Delta t$) and fit each one to the theoretical prediction $P_{\text{fluc}}$. **A)** The inferred parameter combination $\Delta t/\tau$ from the fit increased with the sampling interval, but slower than the expected (linear). The inferred parameters of a few species oscillated strangely, particularly at large $\Delta t$. **B)** The predicted distribution $P_{\text{fluc}}$ changes very little for $\frac{\Delta t}{\tau} \gtrapprox 1$. This makes inferring $\Delta t/\tau$ difficult once $\Delta t$ approaches $\tau$, which is the case for the microbiome. The inset shows the Jensen-Shannon Distances (JSD) between the distributions at the indicated $\Delta t/\tau$ values and $\Delta t/\tau = 40$.The distance between these distributions vanishes quickly once $\Delta t/\tau$ crosses 1. **C-F)** The stiffness of the parameter inference procedure once $\Delta t \approx \tau$ causes the inferred parameter of a species to oscillate due to small changes in the underlying data. The panel shows the data for OTU 56, which had an ENR of 2.47. The ENR in panel B was 2.5. Fits of the LFD were performed at a fixed ENR, which was obtained from fitting the abundance distribution.

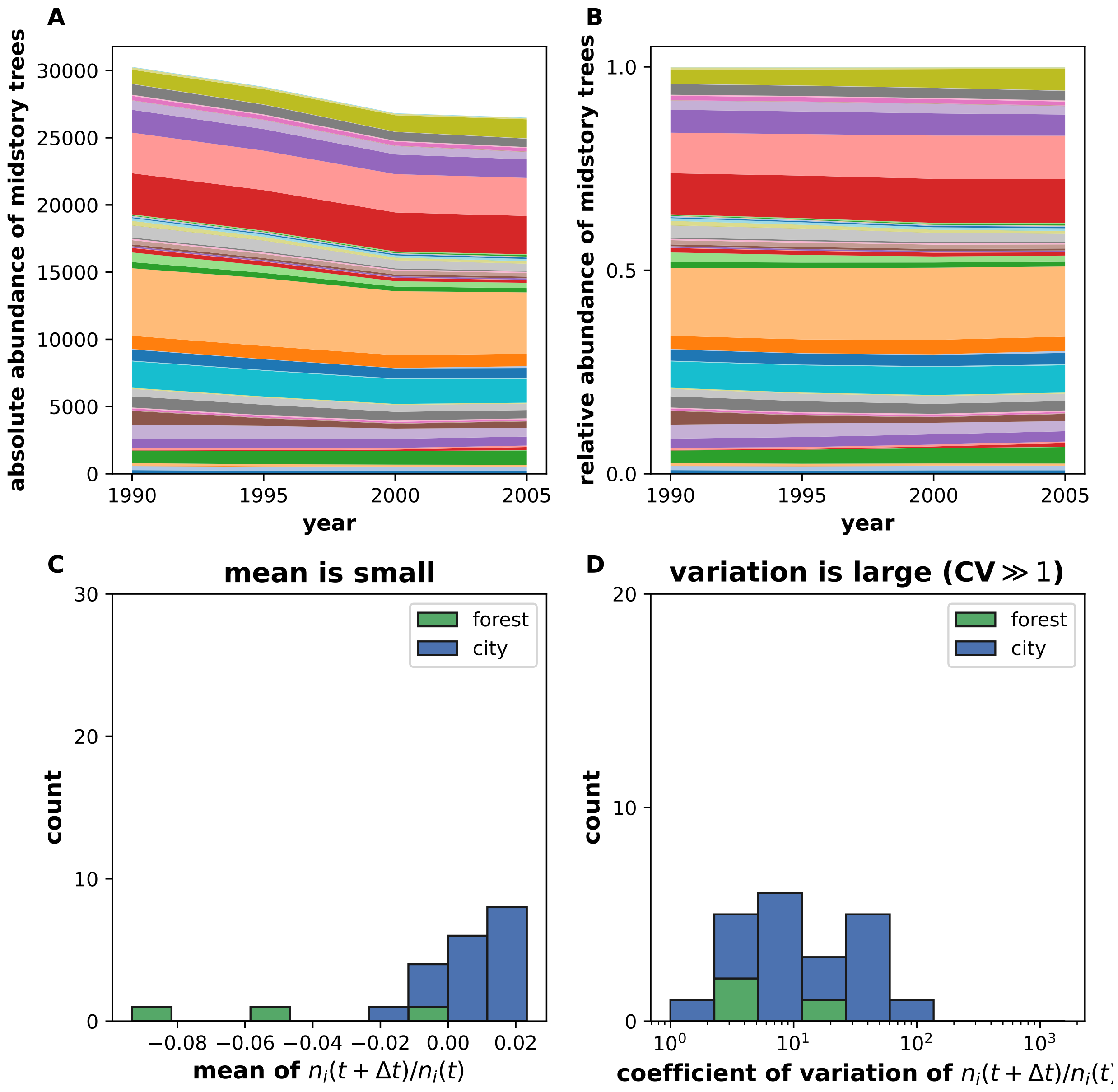

Figure S10: **The temporal trends in absolute population sizes and the importance of fluctuations in logarithmic fold-change of absolute abundance data.** Absolute abundance data may display apparent trends over observation time scales. A) The total population sizes of tree clusters, such as midstory trees (shown), decreased over the observation time window. B) Studying the relative abundances allows us to disentangle the effects of changes in overall population sizes from the fluctuations of species within the overall population cluster. Colors denote different species within the cluster. C) The mean logarithmic fold-change computed from absolute abundance data, $\frac{n_i(t+\Delta t)}{n_i(t)}$, of each trajectory is smaller than the logarithmic fold-change values in Fig. S11. D) The coefficient of variation (ratio of standard deviation to mean) is larger than one in both data sets. This allows us to neglect the small change in the mean in comparison to the large fluctuations between consecutive time points, and fit our model to the empirical LFD in Fig. S11. Both panels show stacked histograms.

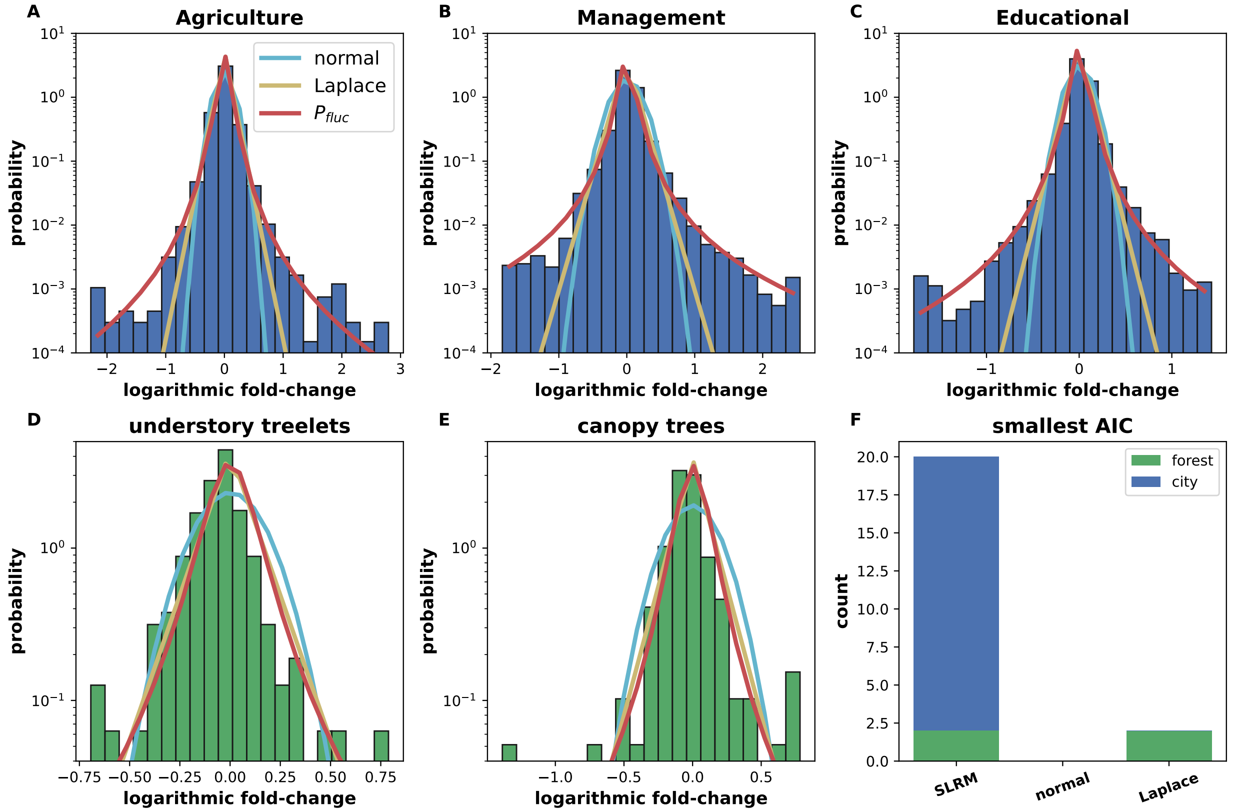

Figure S11: **Fitting the LFD obtained from count data.** The empirical Logarithmic Fold-change Distribution (LFD) is computed using absolute abundances instead of relative abundances for cities and forests, where such data is available. The empirical LFD is shown for three employment sectors **(A-C)** and two tree height clusters (**(G,H)**. The LFD is fit by three candidate distributions: normal(cyan), Laplace (yellow), and the prediction from the SLRM, $P_{\text{fluc}}$ (red). **(I)**The SLRM prediction has the smallest Akaike Information Criterion [38] value in all of the employment data and 2 of the 4 forest clusters. When fitting LFD from absolute abundance data, temporal fluctuations in overall population size can contribute if they are large enough. This may explain why SLRM performed better than the other candidate distributions in all four forest clusters for relative abundances but only two clusters for absolute abundances.

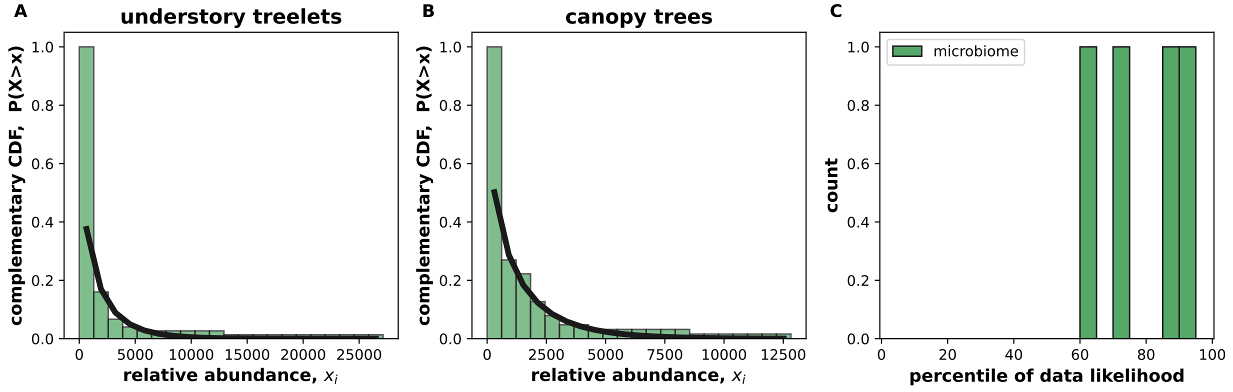

Figure S12: **Fitting the SSD obtained from count data.** The empirical SSD (LFD) is computed using absolute abundances for forests. **A,B)**The empirical distribution of abundances of two tree height clusters and the fit by the model. **C)** All of the fits pass the likelihood-based goodness of fit test. The same analysis cannot be done for cities since we are considering different cities, with different total population sizes

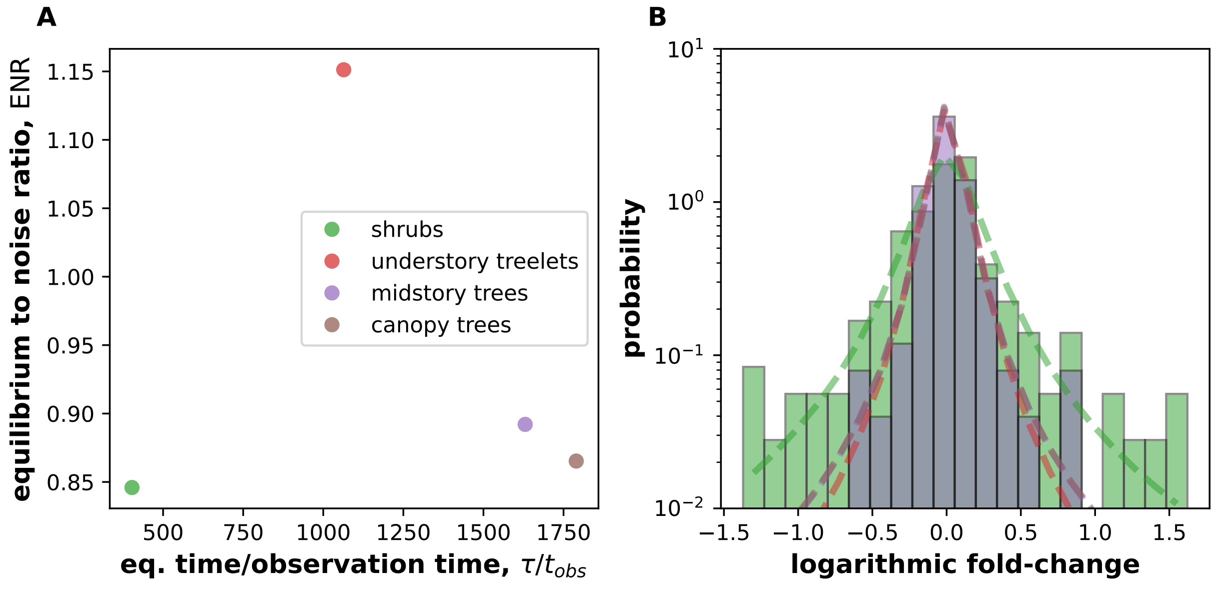

Figure S13: **Comparing forest clusters. A)** The ENR and equilibration timescale $\tau$ obtained by fitting the LFD of the four tree clusters. Shrubs have a much shorter equilibration timescale than the other three clusters. The ENR varies over only a small range of values. **B)** The predicted LFD distributions at the best-fit parameters are plotted, along with the underlying data (only two of the four clusters shown for clarity). The LFD of shrubs differs significantly from the LFDs of understory treelets, midstory trees, and canopy trees, which are all similar to each other. Notably, shrubs experience many more large fluctuations than the other three categories.
